# Optogenetic hedonic hotspots in orbitofrontal cortex and insula: causing enhancement of sweetness ‘liking’

**DOI:** 10.1101/2024.07.31.606067

**Authors:** Ileana Morales, Kent C. Berridge

## Abstract

Hedonic hotspots are brain subregions that causally amplify the hedonic impact of palatable tastes, measured as increases in affective orofacial ‘liking’ reactions to sweetness. Previously, two cortical hedonic hotspots in orbitofrontal cortex and insula were identified by neurochemical stimulation using opioid or orexin microinjections. Here we used optogenetic stimulation in rats as an independent neurobiological technique for activating cortical hedonic hotspots to identify hedonic functions and map boundaries. We report that channelrhodopsin stimulations within rostral orbitofrontal and caudal insula hotspots doubled the number of hedonic ‘liking’ reactions elicited by sucrose taste. This independently confirms their robust functional identity as causal amplifiers of hedonic ‘liking’ and confirms their anatomical boundaries. Additionally, we confirmed an intervening suppressive hedonic coldstrip, to stretching from caudal orbitofrontal cortex to rostral insula. By contrast to localized hedonic hotspots for ‘liking’ enhancement, motivational ‘wanting’ for reward, measured as laser self-stimulation, was mediated by more widely distributed anatomical sites.

## Introduction

Reward contains multiple core components of ‘liking’, ‘wanting’, and learning. Among those, possibly the least understood remains the neural mechanisms of ‘liking’, namely, the immediate hedonic impact of a pleasant stimulus. However, progress has been gained by identification of ‘hedonic hotspots’, or small subregions within mesocorticolimbic brain structures that are uniquely capable of enhancing ‘liking’ reactions to sweetness when neurochemically stimulated, based on affective taste reactivity studies in rats (*1–6*).

Hedonic hotspots were originally identified in subcortical structures, such as nucleus accumbens shell (NAc), ventral pallidum (VP), and brainstem pons (1, 2, 6). More recently two hedonic hotspots were also identified in cortex: an 8 mm^3^ subregion of rostromedial orbitofrontal cortex (OFC), and a 6 mm^3^ subregion of posterior insula cortex (7). Each of those studies used local microinjections in rat hotspots of either opioid, orexin, or endocannabinoid agonists to double or triple affective orofacial expressions of positive ‘liking’ that are elicited by sweetness and other pleasant tastes, versus negative ‘disgust’ elicited by bitterness and other unpleasant tastes, in human infants, other primates, and rats (8–10).

Opioid/orexin stimulation of the OFC hotspot that increased sucrose ‘liking’ also recruited Fos increases in the insula hedonic hotspot, as well as vice versa (7). Either OFC or insula stimulation further increased Fos in other subcortical hedonic hotspots in rostrodorsal NAc medial shell and posteriolateral VP. That supported the hypothesis that ‘liking’ enhancement induced by neurochemical stimulation of any one hotspot requires recruitment of other hotspots in simultaneous neural co-activation as a unified hedonic network to increase hedonic impact (7, 11). Between the rostral OFC and caudal insula hotspot, an 18 mm^3^ suppressive hedonic ‘coldstrip’ was also identified, comprising posteriolateral OFC and rostral insula, where the same orexin or opioid microinjections oppositely reduced ‘liking’ reactions to sucrose taste and failed to recruit Fos activation in other hedonic hotspots.

However, the exclusive use of pharmacological stimulations in hedonic hotspots to enhance ‘liking’ reactions raises the question of whether hedonic hotspots are mere neurochemical artifacts, limited to the effects of local drug microinjections or instead are robust neurofunctional entities for enhancing ‘liking’? If the latter, hedonic hotspot capacities should also be able to confirmed, at least in principle, by triangulating evidence from independent nonpharmacological neural stimulations that similarly enhance ‘liking’ reactions. To potentially provide such independent confirmation that hedonic hotspots have special capability for ‘liking’ enhancement, here we assessed whether optogenetic channelrhodopsin (ChR2) stimulations in cortical hotspots would alter ‘liking’ reactions to sucrose or quinine tastes. In support of this optogenetic effort, we noted that others have reported previous optogenetic studies indicating that ChR2 stimulation in the anterior insula of mice elicited positive affective taste-elicited expressions (12), and that optogenetic stimulation in a gustatory region of insula promoted intake of palatable solutions and ingestive patterns of spout licking in mice (*13*, *14*).

We assessed further whether 1) optogenetic stimulation in previously identified orbitofrontal or insula hotspots enhanced positive ‘liking’ reactions to sweetness, 2) the anatomical locations and boundaries of cortical hedonic hotspots when mapped optogenetically were similar to those previously mapped by neurochemical stimulations (*7*), and 3) whether a suppressive hedonic coldstrip intervened between OFC and insula hotspots, where optogenetic stimulations reduced ‘liking’ reactions to sweetness. Finally, using laser self-stimulation tests, we assessed whether sites able to support motivational ‘wanting’ to obtain an incentive were more widely distributed across cortex, extending outside of hedonic hotspots (*7*). Our results suggest that neurochemically mapped OFC and insula hotspots are indeed able to enhance hedonic ‘liking’ reactions to sweetness when activated optogenetically, with similar anatomical boundaries, and that a cortical optogenetic hedonic coldstrip also exists between them.

## Results

### Local Fos and Neuronal Spread of Activation

Fos expressing cells around optic fibers were counted to measure local ‘Fos plumes’ induced by ChR2 laser stimulation (**Fig. 1, A to C**). The averaged diameters of Fos plumes were used to set symbol sizes in maps showing localization of function of hedonic enhancement sites or hedonic suppression sites based on taste reactivity data and laser self-stimulation data. Localization of function maps were used to map the anatomical boundaries and calculate volumes of optogenetic hedonic hotspots and suppressive coldspots in OFC and insula. Fos plumes typically had a 2-layer structure, with inner zones of intense 250% Fos elevation averaging 0.54 ± 0.05 mm in diameter (volume = 0.08 mm^3^) surrounded by outer plumes of moderate 150%-250% Fos elevation averaging 0.9± 0.04 mm in diameter (volume =0.39 mm^3^), compared to control baseline Fos levels measured at equivalent sites in laser-illuminated eYFP rats (**Fig. 1, A to D**) and also relative to unoperated naïve control rats (**Fig. S1, A to C**). The size of inner plumes (F3,25 = 2.06, *p <* 0.13) and outer plumes did not differ across cortical sites (*F*3,25 = 0.98, *p <* 0.42; *n* = 11 rostral OFC, *n =* 9 caudal OFC, *n* = 4 caudal insula, *n =* 5 rostral insula), and so the outer plume diameter of 0.90 mm was used to set the maximum size of individual site symbols in anatomical localization of function maps. The color of symbols in those maps represents intensity of functional effects induced by ChR2 stimulation at that site: optogenetic-induced changes in hedonic ‘liking’ reactions to sucrose, or of optogenetic self-stimulation, expressed as a within-subject ChR2-induced percent change compared to control no-laser baseline measured in the same rat.

**Fig. 1.**
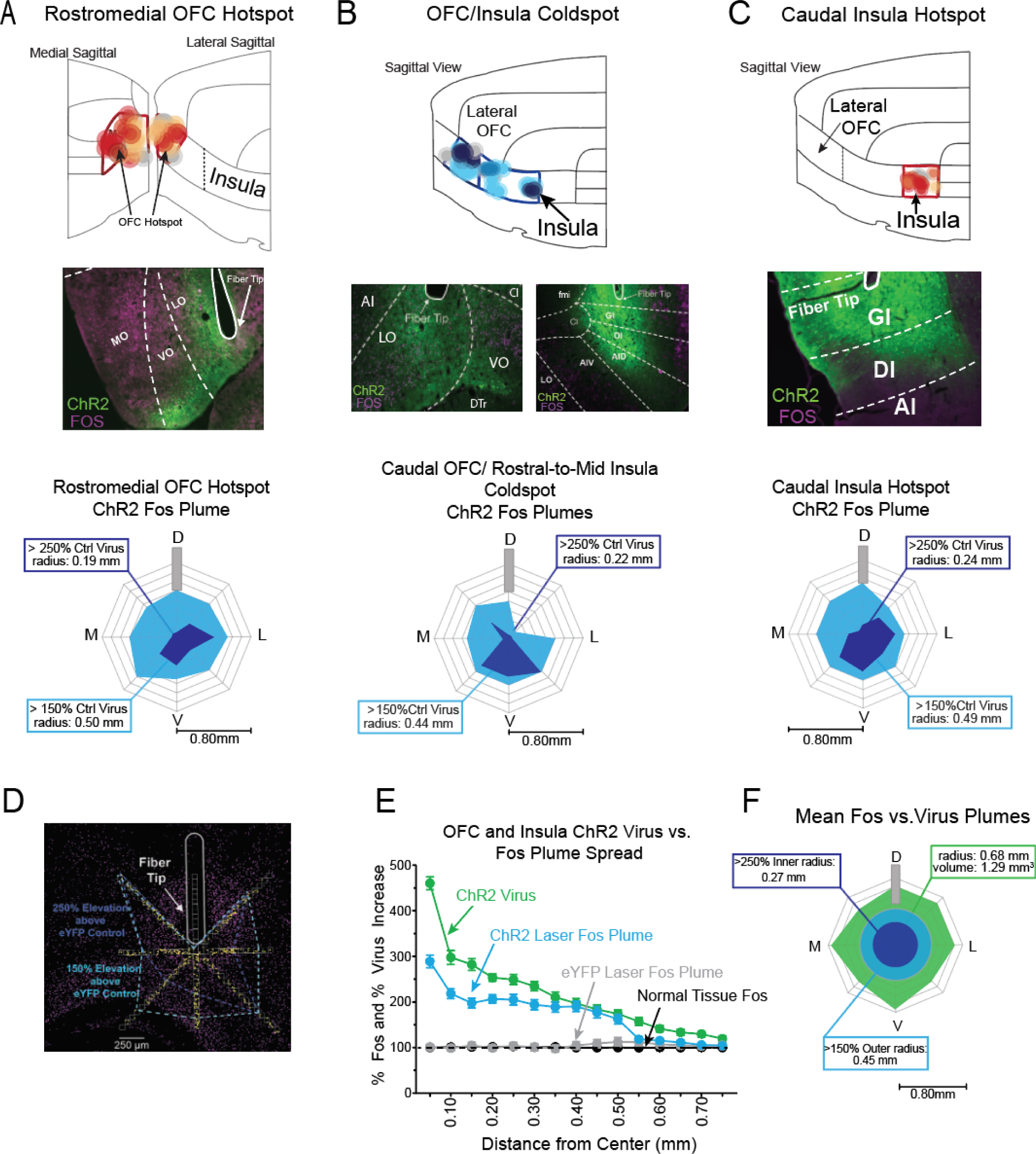
Cortical hotspot ChR2 virus and Fos plumes. (**A**) Rostromedial OFC sites of hedonic enhancement (medial and lateral sagittal views). Rostromedial OFC photomicrograph (10x magnification) shows green channelrhodopsin (ChR2) virus infection (AAV5-ChR2-eYFP) and magenta Fos protein. Average ChR2 laser-induced Fos plume in rostromedial OFC (>250% above eYFP: light solid blue, > 150% above eYFP rats: dark solid blue (**B**) Caudolateral OFC and rostral insula sites of hedonic suppression in OFC/insula coldstrip (lateral sagittal view). Caudolateral OFC and rostral insula photomicrographs showing green channelrhodopsin (ChR2) virus infection (AAV5-ChR2-eYFP) and magenta Fos protein. Average ChR2 laser-induced Fos plume in caudolateral OFC and anterior insula- (>250% above eYFP: light solid blue, > 150% above eYFP rats: dark solid blue. (**C**) Caudal insula sites of hedonic enhancement (lateral sagittal view). Caudal insula photomicrograph showing green channelrhodopsin (ChR2) virus infection (AAV5-ChR2-eYFP) and magenta Fos protein. Average ChR2 laser-induced Fos plume in caudal insula (>250% above eYFP: light solid blue, > 150% above eYFP rats: dark solid blue. (**D**) Photomicrograph shows Fos expression example of local fos plume surrounding fiber in the OFC hotspot. (**E**) Left graph shows how quantitative increases in virus and in Fos protein decline as a function of distance from the fiber tip (Combined rostromedial OFC virus, caudolateral OFC and anterior insula virus, and caudal insula virus, n =26, Combined rostromedial OFC laser Fos, caudal OFC/ anterior insula laser Fos, and caudal insula laser Fos, n =21; Ctrl eYFP Fos, n = 19; Naïve tissue Fos, n = 6). All data represented as mean and standard error (SEM). Right shows average ChR2 virus spread away from fiber optic tip relative to average size of Fos plume. D: dorsal, M: medial, V: ventral, L: lateral.

Fos plumes were typically smaller than the zone of ChR2/eYFP virus expression (**Fig. 1, E,F**), suggesting there was a minimum threshold of laser stimulation required to elevate Fos excitation in ChR2 infected neurons. The mean ∼1.4 mm diameter (1.3 mm^3^ volume) of virus infection did not differ between OFC and insula sites (F3,22 = 0.93, *p <* 0.44; *n* = 9 rostral OFC hotspot, *n* = 4 caudal insula hotspot, *n =* 9 caudal OFC coldspot, *n =* 4 rostral to mid insula coldspot). Since Fos plumes were smaller than zones of ChR2 virus infection (**Fig. 1E**), plume diameter was taken as the best indicator of how far neuronal excitation spread from an optic fiber tip.

### Hedonic taste reactivity

*Baseline affective reactions elicited by tastes.* Oral infusions of sucrose solution at both 0.03M and 0.1M concentrations elicited positive ‘liking’ reactions in both female and male rats on control baseline trials without laser. Dilute 0.03 M sucrose (**Fig. S2, A**) elicited moderate numbers of positive reactions (e.g., lateral tongue protrusions; rhythmic midline tongue protrusions; paw licking; *M* =12.24, *SEM* = 0.65; *F*1,79 = 167.0, *p* < 0.0001, *n =* 81). Very few negative aversive reactions were elicited by 0.03M sucrose (e.g., gapes, headshakes, forelimb flails, etc; *M* = 3.06, *SEM* = 0.38). A baseline sex difference was found to the low sucrose concentration, as females emitted about 20% more ‘liking’ reactions to dilute 0.03M sucrose than males in absence of laser ((*F*1,79 = 6.28, *p* =0.01, female ‘liking’: *M* = 13.56, *SEM* = 0.86; *n =* 40 females, male ‘liking’ score: *M* = 10.95, *SEM* = 0.98, *n* = 41; *t*158 =2.66, *p* = 0.02; 95% CI = [0.4, 5.0], *d* = 2.9). However, more concentrated 0.10 M sucrose (**Fig. S2, B**) elicited higher numbers of positive ‘liking’ reactions *(M* = 17.19, *SEM* = 1.12; *F1,38* = 163.9, *p* < 0.0001, *n = 40)* similarly from both sexes, and hardly any negative ‘disgust’ reactions (*M* = 2.55, *SEM* = 0.37;); Sex x valence: *F*1,38= 0.78, *p* = 0.38, *n* = 20 males, *n* = 20 females).

Conversely, baseline oral infusions of bitter 3x 10^-4^ M quinine in absence of laser (**Fig. S2, C**) solution elicited predominantly negative ‘disgust’ reactions (e.g., gapes, headshakes, forelimb flails; face washing; ‘disgust’ score *M* = 33.03, *SEM* = 1.73; *F1,66* = 308.4, *p* < 0.0001, n = 68), and almost no positive ‘liking’ expressions (‘liking’ *M* = 1.31, *SEM* = 0.24,), with no sex differences detected (*F*1,66= 1.24, *p* = 0.27, *n* =33 females, *n* = 35 males). Finally, oral infusions of water (**Fig. S2, D**) elicited moderate numbers of both positive and negative reactions, although water still elicited slightly more positive ‘liking’ reactions than aversive ‘disgust’ expressions from both male and female rats (‘liking’ reactions, *M* = 9.83, *SEM* = 0.67; ‘disgust’ reactions *M* = 7.14, *SEM* = 0.70, valence*: F1,68* = 7.33, *p* = 0.001, *n* = 70). Again, females emitted slightly more positive ‘liking’ reactions to water than male rats in the absence of laser stimulation (sex: *F*1,68= 5.78, *p* = 0.02, *n* = 33 females, *n* = 35 males).

### Rostromedial OFC hedonic hotspot: optogenetic ChR2 stimulation enhances ‘liking’ reactions

*OFC ChR2 enhancement of sucrose ‘liking’ reactions*. At ChR2 sites within an anteromedial subregion of OFC (**Fig, 2, A; Fig. S3)**, which was previously identified by opioid/orexin microinjections as a hedonic hotspot (*7*), laser stimulation (5, 10, 20, 40 Hz; 5-s ON/ 5-s OFF) approximately doubled (**Fig. 2, B; Fig. S4, A,B)** the overall number of positive hedonic reactions elicited by oral sucrose infusions of both 0.03M or 0.1M concentrations (rhythmic tongue protrusions, lateral tongue protrusions, and paw licks). Laser illumination increased positive ‘liking’ reactions in anteromedial OFC ChR2 rats by 235% ± 32% for 0.03M sucrose over measured control baseline levels in the same individuals without laser (**Fig. S3, A;** 0.03M Sucrose: laser x valence x virus *F*1,34 = 16.39, *p* = 0.0003; *n* = 23 ChR2; *n* = 13 eYFP; ChR2 Sum of rhythmic tongue protrusions, lateral tongue protrusions, and paw licks: *Laser-ON*: *M* = 16.52, *SEM* = 1.22; *Laser-OFF*: *M* = 8.80, *SEM* = 0.87; paired comparison *t*23 = 8.30, *p* < 0.0001; 95% CI = [-10.1, -5.3] *d* = 7.3).The magnitude of laser enhancement of ‘liking’ reactions was similar in females and males (**Fig. S4, C**; ChR2: sex x valence x laser interaction: *F*1,21 = 0.76 *p* = 0.39; *n* = 12 males *n* = 11 females). Anteromedial OFC ChR2 laser stimulation did not alter the few negative ‘disgust’ reactions elicited by 0.03M sucrose (**Fig. S4, A**; gapes, headshakes, forelimb flails; chin rubs; *Laser-ON*: *M* = 3.24, *SEM* = 0.62, *Laser-OFF*: *M* = 2.30, *SEM* = 0.42, *t*22 = 0.94, *p* = 0.72).

**Fig. 2.**
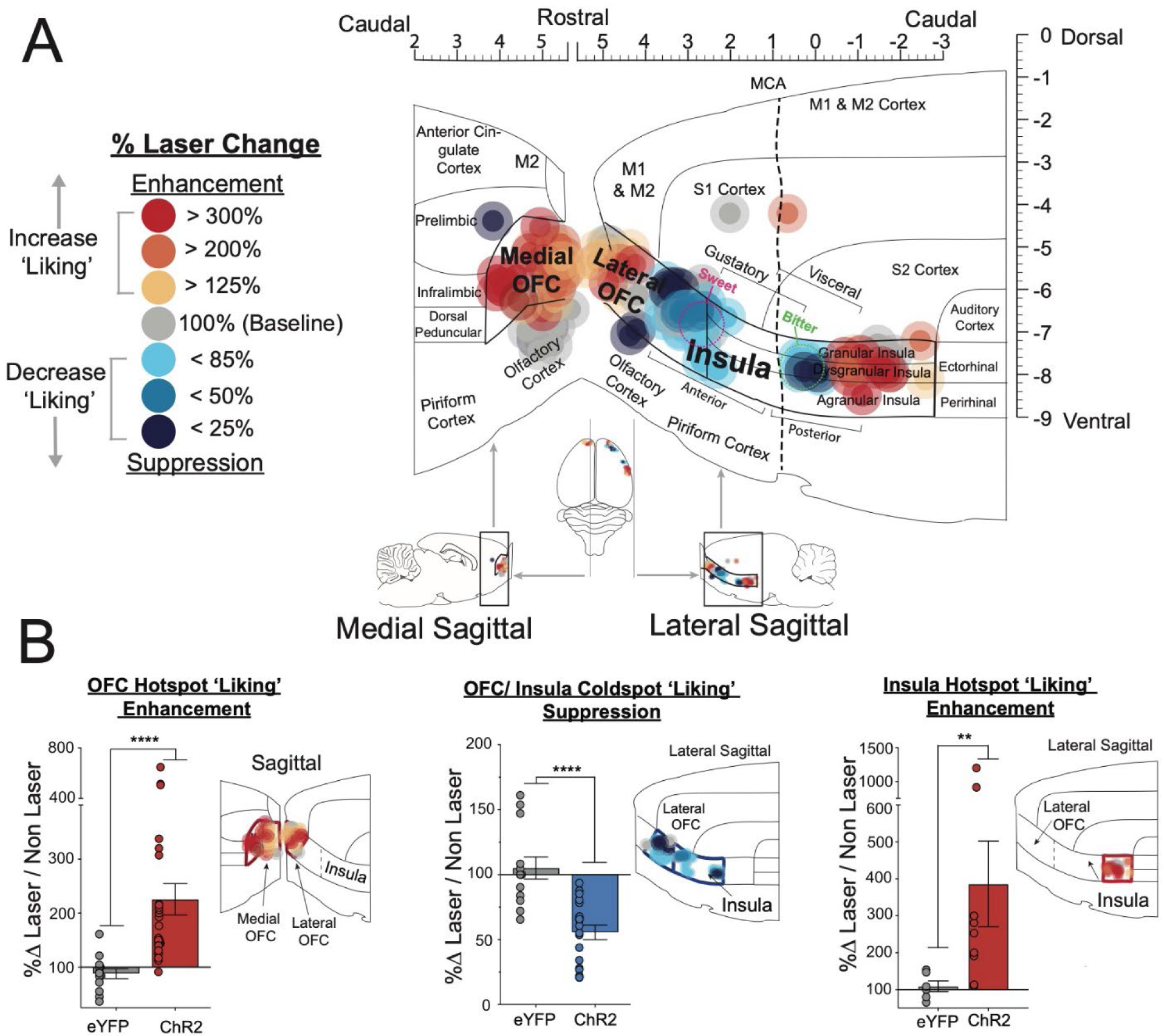
Cortical sites in orbitofrontal cortex and insula that support hedonic enhancement or suppression. (**A**) Localization of hedonic function map shows how optogenetic ChR2 stimulation altered the hedonic impact of sucrose at each individual’s cortical site. Colors reveal hedonic enhancement/suppression effects of ChR2 laser stimulation at each cortical site, measured as laser-induced changes in hedonic taste reactivity (positive ‘liking’ reactions) elicited by intraoral sucrose infusions. Each symbol placement indicates an individual rat’s site size of symbol reflects average size of Fos plumes). Color of symbol represents the within-subject behavioral change in hedonic reactions induced by ChR2 laser stimulation reflected as percent change from no laser control conditions measured in the same rats (‘Liking’ enhancements: red-yellow; ‘Liking’ suppression: Blue). (**B**) Laser ChR2 stimulations differentially alter hedonic ‘liking’ reactions depending on the anatomical subregion of OFC and insula. At rostromedial OFC hotspot and caudal insula hotspot sites, laser stimulation enhanced hedonic ‘liking’ reactions 200% - 300% in ChR2 rats, but not in eYFP controls (rostromedial OFC: U = 18.00, ****p < 0.0001; caudal insula: U = 4.00, **p < 0.01. In the intervening coldstrip, spanning caudolateral OFC to mid insula, laser ChR2 stimulations oppositely suppressed sucrose ‘liking’ reactions to approximately 50% in ChR2 rats, but not in eYFP controls (OFC/insula coldstrip (U = 12.00, ****p < 0.0001)). Data presented as means and standard error (SEM). Anatomical abbreviations: M1: primary motor cortex, M2: secondary motor cortex, S1: primary somatosensory cortex. S2: secondary somatosensory cortex. Gustatory insula zones adapted from (79), visceral insula functional zone adapted from (80) and sweet/bitter coding regions adapted from (13)

Similarly, for the higher sucrose 0.1 M concentration (**Fig. S4, B**), anteromedial OFC laser stimulation nearly doubled (183 ± 21%) the number of positive hedonic reactions compared to within-subject baseline levels (laser x virus x valence: *F1,18= 23.14*, *p* = 0.0001; Laser-ON *M =* 19.36*, SEM =* 2.27*; Laser OFF M* = 11.45, *SEM* = 1.45; ChR2 ‘liking’: *t*36 = 5.4, *p* < 0.0001; 95% CI= [-11.8, -4.1], *n =* 11 ChR2). The percentage magnitude of laser hedonic enhancement of ‘liking’ reactions over baseline levels was comparable for both 0.03M and 0.10 M sucrose (*F1,16=* 0.12, *p* = 0.74), and the magnitude of laser enhancement of hedonic ‘liking’ reactions to 0.1M sucrose (**Fig. S4, D**) was similar in male and female ChR2 rats (*F*1,9 = 0.46 *p* = 0.52; *n* = 4 males *n* = 7 females)

By contrast, in control eYFP rats (**Fig. S4, A, B**) with optically inactive virus, laser illumination in anteromedial OFC sites failed to alter either positive hedonic reactions or negative ‘disgust’ reactions to sucrose from baselines measured in the same individuals, for either 0.03 or 0.1 M sucrose (0.03M sucrose ‘liking’: *t*34 = 2.0, *p* = 0.20; 0.03M sucrose ‘disgust’: *t*34 = 0.03, *p* > 0.99; *n* = 13; 0.10M sucrose ‘liking’: *t*36 = 2.2, *p* = 0.14; 0.10M sucrose ‘disgust’: *t*36 = 1.9, *p* = 0.29; *n* = 9). No sex differences (**Fig. S4, E**) in affective reactions were detected between this group of male and female rats (sex x valence x laser interaction: 0.03M sucrose: *F*1,11 = 0.19 *p* = 0.67; *n* = 7 males *n* = 6 females; 0.10M sucrose: *F*1,7 = 0.001 *p* = 0.96; *n* = 5 males *n* = 4 females).

*Different laser frequencies produce similar enhancements:* Hedonic enhancement effects were robust and similar across a range of different laser frequencies in the rostromedial OFC hotspot (5, 10, 20, 40 Hz; 1-3 mW intensity), and was not limited to any single frequency. All frequencies produced similar magnitudes of enhancements of positive ‘liking’ reactions to sucrose (**Fig. S4, F**), ranging between ∼150% - 300% above within-subject no-laser baselines, and did not differ statistically from each other (*F3,67 = 1.01, p =* 0.39). Assessed separately, 5 Hz, 10 Hz, 20 Hz and 40 Hz frequencies each increased positive hedonic reactions by 150% - 250% above no-laser baselines measured in the same rats (20 Hz *=*211% ± 24% increase, *F1,19=* 40.54, *p* < 0.0001; ‘liking’: *t*19 = 8.41, *p* < 0.0001, 95% CI= [5.5, 10.0], *d=* 9.8, *n = 20;* 10 Hz =240% ± 59 % increase, *F1,13=* 12.01, *p* = 0.004, *n = 14*, ‘liking’ paired comparison: *t*13 = 5.5, *p* = 0.0002, 95% CI= [3.8, 10.3], *d=* 4.0, *n =* 14; 5 Hz = 157% ± 16% increase, *F1,13= 14.17*, *p* = 0.0024, ‘liking’ paired comparison: : *t*13 = 5.5, *p* = 0.0002, 95% CI= [3.0, 8.0], *d=* 3.4, *n =* 14).

*Anatomical boundaries of optogenetic hedonic hotspot in anteromedial OFC:* Localization of function was mapped for optogenetic changes in hedonic reactions caused across OFC and insula sites (**Fig. 2A; Fig. S3**). Hedonic enhancement sites were considered to be sites where ChR2 laser illumination caused 150% - 400%+ increases in ‘liking reactions to sucrose, compared to no-laser baseline levels measured in the same individual. Function maps revealed that hedonic enhancement sites clustered anatomically into two cortical hedonic hotspots: anteromedial OFC and far caudal insula (**Fig. 2A; Fig. S3**).

The anterior border of the OFC hedonic hotspot began at the far rostral tip of OFC (∼+5.64 mm AP), near the anterior edge of medioventral orbital cortex, and then extended caudally along both lateral and medial surfaces. The OFC hotspot was bordered dorsally on the medial surface by prelimbic cortex, and dorsally on the lateral surface by secondary motor cortex. Moving posteriorly along the medial surface, the OFC hedonic hotspot extended ∼1.4 mm to the far caudal edge of medial orbital cortex (∼ +4.28 mm AP). Along the lateral surface, the hotspot extended posteriorly ∼2.1mm to a point approximately ∼3.72 mm anterior to Bregma. There the hotspot was bordered dorsally by the claustrum, medially by the dorsal peduncular cortex, and laterally by the rostral insula. Overall, the OFC hedonic hotspot thus extended rostrocaudally (AP) in length ≈ 2.1mm, mediolaterally (ML) ≈ 2.4 mm, and dorsoventrally (DV) ≈ 2.2 mm, with a total volume of ≈ 11.1 mm^3^. We note these optogenetic OFC hotspot boundaries corresponded closely to those originally mapped neurochemically using opioid/orexin microinjections (*7*). The only difference between our optogenetic map and the earlier opioid/orexin map is that our study probed further in a dorsolateral direction than the earlier microinjections, and we found that the optogenetic OFC hotspot additionally extended into the rostral tip of the dorsolateral orbital cortex, making our total volume slightly larger by ∼25%. Thus, portions of medial orbital (MO), ventral orbital (VO), lateral orbital (LO), and dorsolateral orbital (DLO) were all included within the ChR2 hedonic hotspot of OFC.

Beyond these boundaries, laser ChR2 stimulations at sites more caudal or lateral in OFC failed to increase ‘liking’ reactions to sucrose taste, including sites in caudolateral orbitofrontal cortex and caudoventral orbitofrontal cortex. Similarly, medial sites in dorsally neighboring prelimbic cortex, or ventrally in neighboring olfactory bulb (**Fig. S5, A**), failed to enhance ‘liking’ reactions to sucrose (*F1,6 =* 0.04, *p* = 0.84, *n* = 7).

*Microstructure of taste reactivity components fits hedonic enhancement pattern.* To confirm that laser ChR2 stimulation within the OFC hotspot caused hedonic enhancements, rather than a mere sensorimotor reaction, we assessed whether changes in individual taste reactivity components were grouped into larger affective categories of positive ‘liking’ versus negative ‘disgust’. For example, a shared increase among multiple components within the positive hedonic category would be required to be categorized as a hedonic increase in positive ‘liking’ reactions (i.e., shared across rhythmic midline tongue protrusions [TP], lateral tongue protrusions [LTP] and paw licks [PL]), without increases in any component of the negative ‘disgust’ category (gapes [G], headshakes [HS], face washes [FW], forelimb flails [FF], or chin rubs [CR] (*8*).

For sites within the OFC hotspot, ChR2-induced enhancements fit this cross-category pattern (**Fig. S5, B**). Hedonic increases were not dominated by any single taste reactivity component, which if so, might have reflected a simpler motor effect. Rather, increases in ‘liking’ elicited by laser-accompanied sucrose taste were distributed across multiple reaction components within the positive hedonic category: (TP: *Laser-OFF: M=* 4.95 *SEM= 0.66; Laser-ON: M=* 8.86, *SEM= 0.81,* laser main effect*: F1,32 = 31.12, p < 0.0001* ; LTP: *Laser-OFF: M=* 2.22 *SEM= 0.38; Laser-ON: M=* 5.28, *SEM= 0.68,* laser main effect*: F1,32 = 27.02, p < 0.0001;* PL: *Laser-OFF: M=* 2.72 *SEM= 0.40; Laser-ON: M=* 3.58, *SEM= 0.40,* laser main effect*: F1,32 = 4.65, p = 0.04.)*, and without increases in reaction components belonging to the negative ‘disgust’ category.

*No detectable motor effects of laser on spontaneous orofacial reactions.* Similarly, providing evidence against mere motor effects, in the absence of any oral infusions, laser illuminations in OFC ChR2 rats failed to induce detectable orofacial movements at any cortical sites **(Fig. S5, C)** in either positive hedonic reactions (*M* = 0.50, *SEM* = 0.34) or negative ‘disgust’ reactions (*M* = 1.5, *SEM* = 0.85; *F1,5 =* 3.00*, p = 0.15*, *n* = 6).

*OFC hotspot hedonic enhancement of water.* Laser ChR2 excitation in the OFC hotspot similarly increased positive ‘liking’ reactions elicited by oral water infusions **(Fig. S5, D)** by >30% over within-subject baselines (*F1,31 = 7.29*, *p* = 0.011, ChR2 paired comparison: *t31=* 3.22*, p =* 0.0, 95% CI= [-6.2, -0.59], *d* = 2.8, *n =* 20 ChR2), with no change in the low number of aversive ‘disgust’ expressions to water (*t31=* 0.79*, p =* 0.99). In absence of laser, oral infusions of tap water at room temperature elicited only a few positive ‘liking’ reactions (*M* = 8.36, *SEM*= 1.08) and a few aversive ‘disgust’ reactions on baseline tests (*M*= 7.38, *SEM* = 1.35). In eYFP control rats, adding laser illumination to OFC sites did not alter either positive or negative reactions to water compared to baseline (‘liking’ paired comparison: *t31=* 1.26*, p =* 0.87, ‘disgust’ paired comparison: *t31=* 0.90*, p =* 0.99, *n* = 12).

*OFC hotspot suppression of quinine ‘disgust’.* Oral infusions of bitter quinine solution (3x10 ^-4^ M) elicited predominately aversive ‘disgust’ reactions in the absence of laser (*M* = 34.74, *SEM* = 2.98; *F1,20 = 140.0, p < 0.0001*, *n* = 21). Within the OFC hotspot, adding laser stimulation in either ChR2 rats or eYFP rats (40 Hz; 5-s ON/ 5-s OFF) moderately suppressed **(Fig. S5, E)** the number of aversive reactions elicited by quinine by about 20%-30% below no-laser baselines measured in the same rats (ChR2 rats: 35% ± 7% suppression, *M*= 20.90, *SEM* = 2.25, *n* = 21 ChR2; eYFP control rats: 18.4% ± 10% suppression, laser x valence: *F1,32 = 34.17, p <* 0.001). The magnitude of quinine ‘disgust’ suppression was nearly twice as large in ChR2 rats as eYFP rats, although this magnitude difference was not significant (*F1,32 = 0.94, p = 0.34; n* = 13 eYFP), suggesting that light or heat from laser in OFC may partly contribute to reduce ‘disgust’ reactions, independently of ChR2-induced neuronal excitation (*15*, *16*). Multiple components of ‘disgust’ reactions elicited by quinine were suppressed **(Fig. S5, F)** together by OFC hotspot laser, supporting the interpretation that the aversive ‘disgust’ of bitterness was reduced: headshakes (*Laser-OFF*: *M* = 6.31, *SEM* = 0.66; *Laser-ON*: *M* = 3.74, *SEM=* 0.46, *t*20 = 5.18, *p <* 0.0001, 95% CI= [-3.6, -1.5], *d* = 4.5), forelimb flails (*Laser-OFF*: *M* = 18.33, *SEM* = 2.34; *Laser-ON*: *M* = 9.10, *SEM= 1.51*, *t*20 = 5.41, *p* < 0.0001, CI= [-12.8, -5.7], *d* = 4.7), and face washes (*Laser-OFF*: *M* = 2.12, *SEM* = 0.42; *Laser-ON*: *M* = 1.29, *SEM=* 0.30, *t*20 = 2.05, *p* = 0.05, CI= [-1.7, 0.01], *d* = 2.3)

*Optogenetic inhibition of OFC hotspot neurons fails to alter affective reactions.* A separate group of rats received the inhibitory virus iC++ at sites in the rostromedial OFC hotspot (*n*=7 total; 4 females, 3 males). Laser inhibition within the OFC hotspot failed **(Fig. S5, G)** to reduce positive hedonic reactions to 0.03M sucrose (*F*1,6 = 0.01, *p* = 0.92; Sum of rhythmic tongue protrusions, lateral tongue protrusions, and paw licks: *Laser-ON*: *M* = 14.64, *SEM* = 3.14; *Laser-OFF*: *M* = 14.36, *SEM* = 2.83). Similarly, optogenetic inhibition in OFC hotspot failed to reduce or alter the moderate number of ‘liking’ reactions to water (**Fig. S5, H;** *F*1,6 = 1.83, *p* = 0.22; *n* = 4 females, *n* = 3 males; Sum of rhythmic tongue protrusions, lateral tongue protrusions, and paw licks: *Laser-ON*: *M* = 6.50, *SEM* = 1.41; *Laser-OFF*: *M* = 7.36, *SEM* = 1.72). Finally, OFC optogenetic inhibition failed to alter predominantly ‘disgust’ reactions elicited by quinine (**Fig. S5, I**; 216% ± 62% laser enhancement; *F*1,6 = 0.26, *p* = 0.63; *n* = 4 females, *n* = 3 males; Sum of face washes, forelimb flails, headshakes, gapes, and chin rubs: *Laser-ON*: *M* = 17.64, *SEM* = 5.63; *Laser-OFF*: *M* = 15.86, *SEM* = 6.16. These lack of inhibition effects suggests that while neuronal excitations in the OFC hotspot are sufficient to generate gains of hedonic function that increase hedonic ‘liking’ reactions, optogenetic inhibition of neurons in the OFC hotspot does not cause a reciprocal loss of normal hedonic function.

### Optogenetic hedonic coldspot strip spans from caudal OFC through rostral insula: ‘Liking’ suppression

*Anatomical boundaries of OFC/Insula optogenetic hedonic coldspot:* Posterior to the caudal boundary of the anteromedial OFC hotspot on the lateral surface of cortex, a hedonically suppressive ‘coldstrip’ began where ChR2 laser stimulation at posterolateral OFC sites oppositely reduced positive ‘liking’ reactions elicited by both 0.03M and 0.10M sucrose tastes (**Fig. 2, A, B**). This suppressive coldstrip extended ∼ 3 mm posteriorly through entire posterolateral OFC, anterior insula and a middle portion of insula, to end at a point in mid-posterior insula just dorsal to where the anterior commissure crosses the midline (AP coordinates ∼+3.00 mm to ∼-0.12mm bregma). The coldstrip included sites in ventral and lateral orbital subdivisions of caudal OFC, caudal dorsolateral OFC and caudal ventrolateral OFC as well as anterior and middle insula. At its rostral tip, the hedonic coldstrip was bordered dorsally by the claustrum, and medially by the dorsal peduncular cortex, and the rostral portion of the coldstrip spanned mediolaterally through the lateral orbitofrontal cortex and anterior insula. At its caudal end, the suppressive coldstrip was bordered dorsally by secondary somatosensory cortex, ventrally by piriform cortex, and medially by the claustrum. Within the insula, the agranular, dysgranular, and granular horizontal layers were all included.

Laser stimulation (40 Hz) at sites in this OFC-insula coldspot strip of ChR2 rats suppressed positive ‘liking’ reactions to 0.03M or 0.1M sucrose to approximately one-half the baseline levels emitted by the same ChR2 rats when no laser was delivered (**Fig. 2, B; Fig. S6, A, B)** (0.03M sucrose: *Laser-ON*: *M* = 9.33, *SEM* = 0.86; *Laser-OFF*: *M* = 17.05, *SEM* = 1.08; *F1,25 =* 5.45, *p* = 0.03; ChR2 ‘liking’ suppression: *t25=* 6.13, *p* < 0.0001, 95% CI = [4.3, 11.1], *d =* 7.9, *n* = 21 ChR2; 0.1M sucrose: *Laser-ON*: *M* = 9.18, *SEM* = 1.25; *Laser-OFF*: *M* = 20.32, *SEM* = 1.69; *F1,21 =* 15.67, *p* = 0.0007; paired comparison: *t42=* 5.83, *p* < 0.0001, *n* = 17 ChR2). The percentage magnitude of suppression was similar for both 0.03M and 0.1M sucrose concentrations (*F3,69=* 0.60, *p* =0.62). Stimulations in the OFC/insula coldstrip also increased the number of aversive ‘disgust’ reactions (**Fig. S6, A, B)** elicited by both 0.03M and 0.10M sucrose(0.03M sucrose: *Laser-ON*: *M* = 7.57, *SEM* = 1.48; *Laser-OFF*: *M* = 3.71, *SEM* = 0.45; *t15=* 3.06, *p* = 0.02; 0.10 M sucrose: *Laser-ON*: *M* = 8.86, *SEM* = 3.58; *Laser-OFF*: *M* = 2.86,*SEM* = 0.63; *t42=* 2.83, *p* = 0.03).

Hedonic suppression of sucrose ‘liking’ reactions was similarly robust (**Fig. S6, C)** across all laser frequencies tested here (*F3,69=* 0.60, *p* = 0.62), and when assessed separately, 40 Hz, 20 Hz, 10 Hz, and 5 Hz each suppressed positive ‘liking’ reactions to sucrose (40 Hz: 41%± 6% decrease, *F1,20=* 29.93, *p* < 0.0001, *n = 21;* 20 Hz: 34%± 6% decrease, *F1,19=* 16.32, *p* = 0.0007, *n = 20;* 10 Hz: 39%± 8% decrease, *F1,15=* 20.21, *p* = 0.0004, *n = 16;* 5 Hz: 28%± 10% decrease, *F1,14=* 12.16, *p* = 0.004, *n = 15).* Similarly, stimulations in both posterior OFC and anterior insula portions of the coldstrip suppressed sucrose ‘liking’ reactions to similar extents, and also similarly increased ‘disgust’ reactions (**Fig S6, D**; OFC/ Insula ‘liking: *F1,19 =* 1.81, *p* = 0.19; OFC/insula ‘disgust’: *F1,19 =* 0.27, *p* = 0.61).

By comparison, in eYFP control rats (**Fig. S6, A, B)**, laser stimulation of sites in the OFC-insula coldstrip failed to alter either positive or negative affective reactions to sucrose (0.03 M Sucrose: *Laser-ON*: *M* = 12.92, *SEM* = 2.71; *Laser-OFF*: *M* = 11.83, *SEM* = 1.57; eYFP ‘liking’: *t25=* 0.46, *p* = 0.99; eYFP ‘disgust’: *t25=* 1.24, *p* = 0.91, *n* = 6; 0.10 M sucrose: *Laser-ON*: *M* = 20.1, *SEM* = 2.6; *Laser-OFF*: *M* = 19.8, *SEM* = 2.5; eYFP ‘liking’: *t42=* 0.08, *p* = 0.99; eYFP ‘disgust’: *t45=* 0.10, *p* = 0.99, *n* = 6)

*Coldstrip microstructure of taste reactivity: Hedonic suppression pattern*. Multiple components within the positive ‘liking’ category elicited by sucrose were suppressed together by ChR2 laser stimulation at sites in the OFC-insula coldstrip (**Fig. S6, E**)(e.g., midline tongue protrusions) (t*21= 4.91*, p < 0.0001, 95% CI= [-9.8, -3.9], *d* = 7.2), and paw licks (t*21= .45*, *p* = 0.003, 95% CI= [-2.6, -0.6], *d* = 3.5)). This suggests coldstrip ChR2 stimulations suppressed positive hedonic reactions as an entire affective category.

W*ater infusions: hedonic suppression*. Optogenetic stimulation (40 Hz) of ChR2 sites within the hedonic OFC-insula coldstrip similarly decreased positive ‘liking’ reactions in response to oral water infusions to approximately one-half the number elicited on control trials without laser in the same rats (**Fig. S6, F)** (*Laser-ON*: *M* = 7.26, *SEM* = 1.00; *Laser-OFF*: *M* = 13.24, *SEM* = 1.24; laser x valence main interaction: *F1,23 =* 15.57, *p* = 0.0006; *n =* 19 ChR2). However, ChR2 rats did not differ significantly from eYFP rats receiving laser, raising the possibility that heat or light from laser may have also impacted affective reactions independently of opsin effects (*Laser-ON*: *M* = 7.67, *SEM* = 1.02; *Laser-OFF*: *M* = 10.58, *SEM* = 2.28; *F1,23 = 2.12*, *p* = 0.16; *n* = 6 eYFP).

*Quinine infusions: Potential suppression of ‘disgust’ reactions.* Laser stimulation of ChR2 sites in OFC-insula ‘hedonic coldstrip’ similarly suppressed aversive ‘disgust’ reactions to bitter quinine (**Fig. S6, G**), by approximately 30% in ChR2 rats *(Laser-ON: M =* 25.48*, SEM =* 2.45*; Laser-OFF: M =* 32.28*, SEM =* 3.05; *laser x valence: F1,20 =* 9.96*, p =* 0.005, *n* = 20 ChR2). However OFC/insula coldstrip activations in eYFP controls suppressed aversive ‘disgust’ reactions by similar levels 23% ± 13%, and eYFP controls did not differ from ChR2 rats, again suggesting heat from the laser source may have contributed to these effects (*laser x valence x virus: F1,20 =* 0.14*, p =* 0.72, *n* = 6 eYFP). Global suppression of both negative aversive reactions to quinine and positive hedonic reactions to sucrose and water, suggests a general affective suppression of both positive ‘liking’ and negative ‘disgust’. Alternatively, it could reflect a general sensorimotor disruption of orofacial reactions. However, in the absence of any taste infusion, laser illumination in OFC/insula coldstrip ChR2 rats failed to produce any detectable orofacial movements on its own (**Fig. S6, H)**(*F1,13 =* 2.48*, p = 0.14*, *n* = 14). Stimulations at posterolateral OFC sites and rostral-middle insula sites within the coldstrip similarly suppressed quinine ‘disgust’ reactions (**Fig. S6, I;** Supplementary Fig 6h *brain region x laser interaction: F1,18 =* 0.36*, p =* 0.55). Only a few positive ‘liking’ reactions were elicited by quinine, and these were not detectably altered by laser ChR2 excitations at coldstrip sites, perhaps because they were near a floor of zero to begin with *(F1,18 =* 1.50*, p =* 0.24).

### A second optogenetic hotspot in far caudal insula magnifies hedonic ‘liking’ to sucrose

In a far-caudal subregion of insula, a second cortical hedonic hotspot was identified (**Fig. 2; Fig S3**), where laser stimulation of ChR2 sites doubled-to-tripled the number of ‘liking’ reactions to sucrose tastes similarly to the rostromedial OFC hotspot. The insula hedonic hotspot included agranular, dysgranular, and granular zones of the farthest caudal one-third of insula, spanning ∼-2 mm from ∼-0.84 mm AP from bregma to ∼-2.92 mm AP (Fig. 2). The caudal insula hotspot was bordered medially by the claustrum, dorsally by secondary somatosensory cortex, ventrally by piriform cortex, and posteriorly by ectorhinal and perirhinal cortex (at its caudal end where it medially abutted external capsule).

Within this caudal insula hotspot, laser stimulation of ChR2 sites increased hedonic ‘liking’ reactions to 0.03M sucrose taste by over 300% ± 116% over baseline levels elicited from the same rats on no-laser trials (**Fig S7, A;** *Laser OFF=* 8.2 ± 1.48; *Laser ON*: = 19.35 ± 2.47; *F*1,14= 10.03, *p* =0.007; *t*14 = 6.4, *p* < 0.0001, 95% CI = [-16.2, -6.1], *d* = 5.5, *n* = 10 ChR2; *n* = 6 eYFP). All laser frequencies (40 Hz, 20 Hz, 10 Hz, and 5 Hz) produced similar magnitude hedonic enhancements at these ChR2 sites (**Fig S7, B**; *F*4, 31 = 1.65, *p* = 0.20), and male and female ChR2 rats showed similar hedonic enhancements in percentage terms, without detectable sex differences (**Fig S7, C***; F*1,8= 0.28, *p* = 0.61; *n* = 4 males, *n* = 5 females). By contrast, in eYFP control rats, laser illuminations in far-caudal insula did not alter either positive ‘liking’ or aversive ‘disgust’ reactions to sucrose (‘liking’: *t*14 = 0.48, *p* > 0.99, ‘disgust’: *t*14 = 1.25, *p* = 0.92).

As caveat, we tested 0.03 M sucrose in all insula hotspot rats but were able to test 0.1M sucrose in only a few rats. That was because we observed laser-induced seizures appear in 50-80% of rats at posterior insula sites after multiple optogenetic ChR2 stimulations, and so restricted most subsequent rats to as few laser stimulations as possible. In the two-posterior insula ChR2 rats we were able to test with 0.10M sucrose, we observed 150% and 133% increases in hedonic ‘liking’ reactions over their no-laser control baselines.

*Water infusions: Hedonic enhancement*. Laser stimulation of ChR2 sites in the far-posterior insula hotspot similarly increased positive ‘liking’ reactions by 500% elicited by water over normally low baselines in the absence of laser (**Fig S7, D;** *F*1,12= 8.24, *p* =0.01; *n* = 9 ChR2) and did not alter affective reactions in eYFP controls (*t*12 = 1.38, *p =* 0.77, *n* = 5). There was no increase in negative ‘disgust’ reactions.

*Taste reactivity microstructure: Hedonic enhancement pattern*. Multiple orofacial components within the positive ‘liking’ category for sucrose and water were increased together by ChR2 laser stimulations in the insula hotspot: rhythmic midline tongue protrusions (*Laser-ON: M = 7.45, SEM = 3.50; Laser-OFF; M = 3.50, SEM 0.84; t*10 = 3.800, *p* = 0.00), and lateral tongue protrusions (*Laser-ON: M = 8.15, SEM = 1.36; Laser-OFF; M = 1.95, SEM 0.50*; *t*10 = 4.21, *p =* 0.002). However, in the absence of any taste infusion, ChR2 laser stimulations failed to elicit any detectable orofacial reactions (positive *‘liking’, M =* 0.5, *SEM =* 0.19; aversive ‘disgust’ *M =* 1.6, *SEM =* 0.42; *F*1,7= 2.39, *p* = 0.17, *n* = 8). This pattern suggests that ChR2 excitation in the posterior insula hotspot specifically enhanced the hedonic impact of tastes that were initially pleasant or neutral, rather than triggering hedonic reactions as a direct effect or motor effect.

*Quinine infusions: no detectable change*. For bitter quinine infusions, laser ChR2 stimulation in the insula hotspot failed to suppress the substantial level of ‘disgust’ reactions, or to increase positive ‘liking’ reactions above their low baselines to bitterness (**Fig S7, F**; *F*1,7= .01, *p* =0.91; *n* = 8). This suggests that a strongly disgusting taste may resist hedonic enhancement by far-posterior insula stimulation.

### Incentive value of laser by itself? Self-stimulation measures

The incentive motivation value of laser ChR2 stimulations on its own, in the absence of any taste infusion, was measured in two laser self-stimulation tests: an active instrumental spout-touch task and a relative passive place-based self-stimulation task.

### Spout-Based Self-Stimulation

*OFC and insula hotspots support active-touch self-stimulation.* In the active-touch self-stimulation task, each instrumental touch on a designated empty waterspout (laser-spout) earned a brief laser bin of either 1-sec or 5-sec duration (depending on test day). By contrast, touching an alternative control spout produced nothing, and served merely as a measure of baseline exploratory touches. Rats were considered to be ‘high self-stimulators’ if they earned >50 laser illuminations in a 30-min session and made at least twice as many contacts on their laser-spout than on control spout (**Fig 3**). Rats were considered to be ‘moderate self-stimulators’ if they earned >10 but <50 illuminations per session, and still made twice as many contacts on laser-spout than on control spout. Finally, rats were considered ‘failures to self-stimulate’ if they earned <10 illuminations or failed to touch the laser spout at least twice as frequently as the control spout. All rats were categorized on day 1 and retested for reliability on days 2 and 3.

**Fig. 3.**
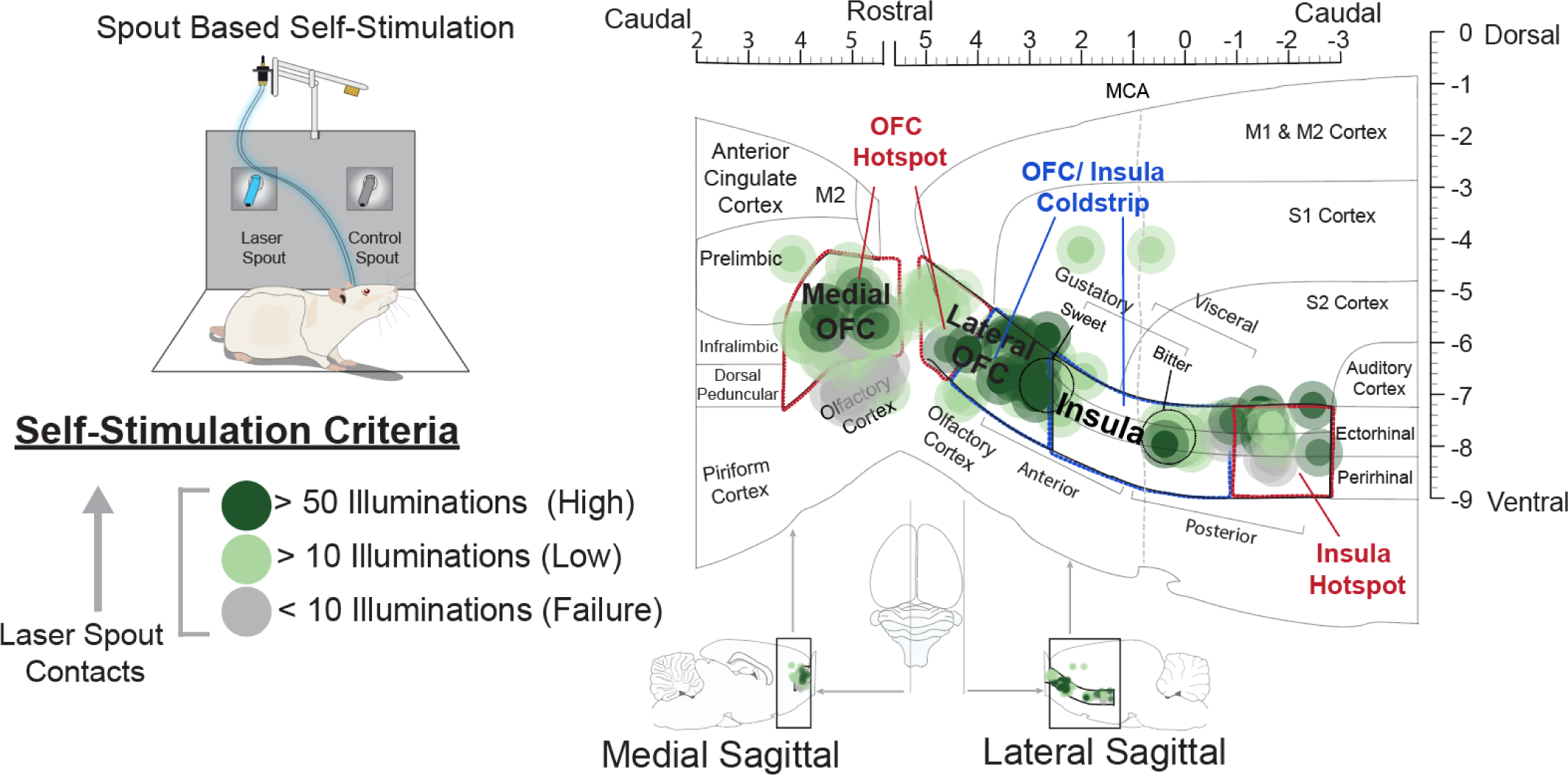
Cortical sites in orbitofrontal cortex and insula that support incentive motivation for reward: laser self-stimulation on spout-touch task. Optogenetic ChR2 stimulation at various cortical sites, both in and outside of hedonic hotspots, support laser self-stimulation. Functional maps show instrumental performance to earn ChR2 laser stimulations at each cortical site on a spout-touch laser self-stimulation task (map based on 40 Hz, 1-s pulse data). Each symbol placement indicates an individual rat’s channelrhodopsin expression (size of symbol reflects size of Fos plumes). Color of symbols represents the level of self-stimulation criteria met by each rat (high self-stimulation (>50 illuminations earned): dark green; low self-stimulation (10 to 49 illuminations earned: light green; Failures to self-stimulate (<10 illuminations earned): grey). For comparison purposes to hedonic ‘liking’ effects, red and blue outlines indicate the anatomical boundaries of the cortical hedonic hotspots and coldstrip mapped based on taste reactivity results in the same rats. Anatomical abbreviations: M1: primary motor cortex, M2: secondary motor cortex, S1: primary somatosensory cortex. S2: secondary somatosensory cortex. Gustatory insula zones adapted from (79), visceral insula functional zone adapted from (80) and sweet/bitter coding regions adapted from (13)

When OFC or insula hedonic hotspot rats could earn brief 1-s 40 Hz laser pulses, ∼77% of OFC hotspot rats and ∼75% of insula hotspot rats met criteria for at least moderate levels of self-stimulation behavior, and 10% to 25% met criteria for high self-stimulation (OFC hotspot: high self-stimulation: 9.1%, moderate self-stimulation: 68.2%, no self-stimulation: 22.7%; Insula hotspot: high self-stimulation: 25.0%, moderate self-stimulation: 50.0%, no self-stimulation: 25%). When touches earned longer 5-s laser pulses (which had been used in taste reactivity tests to increase hedonic ‘liking’ reactions to sucrose) ∼ 50% to 60% met criteria for at least at moderate -levels of self-stimulation, and 10% - 20% met criteria for high self-stimulation (5-sec OFC hotspot: high self-stimulation: 18.2%, moderate self-stimulation: 45.5%, no self-stimulation: 36.4%; Insula hotspot: (high self-stimulation: 12.5%, moderate self-stimulation: 37.5%, no self-stimulation: 50.0%). OFC hotspot sites were slightly more effective than insula hotspot sites at promoting self-stimulation when longer 5-s pulses were delivered (**Fig. S8, A**; 20 Hz laser x brain region x pulse duration interaction: *F*1,19= 8.54, *p* = 0.0009; OFC 1-s: *t*19 = 3.1, *p =* 0.02, 95% CI= [-25.9, -1.7], *d* = 4.1; OFC 5-s: *t*19 = 3.5, *p =* 0.01, 95% CI= [-26.5, -3.0], *d* = 3.5; Insula 1-s: *t*19 = 2.8, *p =* 0.05, 95% CI= [-37.6, 0.09], *d* = 3.1; 40 Hz laser x brain region x pulse duration interaction: *F*1,28= 3.56, *p* = 0.07).

By contrast, eYFP rats with sites in OFC or insula hotspots failed to meet any criteria for laser self-stimulation in the spout-touch task. Control eYFP rats made similar numbers of contacts on the laser spout and non-laser spout, both for 1-s laser bins (**Fig. S8, A**; Spout contacts: Laser-ON: M =17.1, SEM = 4.6; Laser-OFF: M = 24.0, SEM = 8.1), and for 5s laser bins (Spout contacts: Laser-ON: M =7.9, SEM = 2.3; Laser-OFF: M = 8.3, SEM = 2.1; laser x pulse duration interaction: *F1,8*= 1.2, *p* = 0.31, *n* = 11).

*Hedonic coldstrip sites also support active-touch laser-self-stimulation.* Many sites in the hedonic-suppressive coldstrip from caudolateral OFC to mid insula also supported laser self-stimulation in the active-touch task, despite having suppressed hedonic ‘liking’ reactions in taste-reactivity tests (**Fig 3, Fig. S8, B;** 20 Hz laser main effect: *F1,18*= 5.55, *p* = 0.03; *n* = 9 OFC, *n* = 11 insula; 40 Hz laser main effect: *F1,18*= 11.87, *p* = 0.003; *n* = 9 OFC, *n* = 11 insula;). When earning 1-sec laser bins, virtually all posterior OFC coldstrip ChR2 rats met criteria for at least low levels of self-stimulation, and ∼50% met criteria for high self-stimulation (caudolateral OFC: high self-stimulation: 55.6%; low self-stimulation: 44.4%; no self-stimulation: 0%). Similarly, ∼70% of insula coldstrip ChR2 rats met criteria for at least low self-stimulation, and ∼25% met criteria for high self-stimulation (anterior & middle insula: high self-stimulation 18.2%; low self-stimulation 54.5%; no self-stimulation 27.3%).

When laser duration was extended to longer 5-s bins, more similar to durations used in the taste reactivity test, coldstrip sites continued to support laser self-stimulation (caudolateral OFC: high self-stimulation: 55.6%; moderate self-stimulation: 33.3%; no self-stimulation 11.1%; Anterior & middle insula: high self-stimulation: 18.2%; moderate self-stimulation: 72.7%; no self-stimulation: 9.1%). Coldstrip sites in both caudolateral OFC and anterior insula supported laser self-stimulation equally at both 1-s and 5-s laser durations, and at both 20 Hz and 40 Hz frequencies (**Fig. S8, B**; 20 Hz laser x brain region x pulse duration interaction: *F1,18*= 0.17, *p* = 0.68; 40 Hz laser x brain region x pulse duration interaction: *F1,18*= 0.18, *p* = 0.68).

These results suggest that stimulations of ChR2 sites in the hedonic coldstrip generate robust incentive value or ‘wanting’ of when tested in the active-touch laser self-stimulation task, despite lack of ‘liking’ enhancement and even suppression of ‘liking’ reactions in the hedonic taste reactivity test. By contrast, eYFP control rats failed to reach any self-stimulation criteria in the spout touch task, and made equal numbers of touches on the laser-delivering spout and non-laser spout for 1-s bins (40 Hz: Spout contacts: *Laser-ON: M* =10.5, *SEM* = 5.7; *Laser-OFF: M* = 31.9, *SEM* = 12.2) and 5s bins (Spout contacts: *Laser-ON: M* =26.8, *SEM* = 17.; *Laser-OFF: M* = 27.3, *SEM* = 13.8; laser x pulse duration interaction: *F1,4*= 2.98, *p* = 0.16, n = 5; 20 Hz laser x pulse duration interaction: *F1,5*= 0.04, *p* = 0.85, *n* = 6).

*Coldstrip vs* O*FC/Insula hotspots: Comparing active-touch self-stimulation.* We explicitly compared laser self-stimulation levels by ChR2 rats with sites in the coldstrip versus sites in either OFC hotspot or insula hotspot (both hotspots combined; n = 30 hotspot rats, n = 20 coldstrip rats). ChR2 rats with sites in the coldstrip earned ∼3x more laser illuminations overall than those with sites in the two hotspots, at both 1-s and 5-s laser bin durations (F 1,48 = 7.71, p = 0.008, ; 1-s bin coldstrip: 105.2 ± 50.4 illuminations; 1-s bin hotspots: 30.6 ± 4.9 illuminations; 5-s bin coldstrip : 71.1 ± 22.7 illuminations; 5-s hotspot rats: 21.9 ± 4.6 illuminations). Thus, if anything, rats with coldstrip ChR2 sites showed greater ‘wanting’ to earn brief laser excitations than rats with hotspot ChR2 sites.

### Place-Based Self-Stimulation Task

*OFC and Insula Hotspots Support Place-Based Self-Stimulation.* In a place-based laser self-stimulation task, rats could earn laser illuminations by entering a designated chamber in a 2-chamber apparatus, or simply remaining in the designated chamber while laser continued to cycle (3 sec ON, 8 sec OFF cycle). Hedonic hotspot sites in both anteromedial OFC and far-caudal insula supported place-based self-stimulation (**Fig. 4, A, B**), as ChR2 rats with OFC or insula hedonic hotspot sites spent 150% - 200% more time in the laser-delivering chamber at both 20 Hz and 40 Hz frequencies than in the alternative chamber without laser (laser main effect; *F*2,73= 4.80, *p* =0.01; *n* = 22 OFC ChR2, *n* = 10 insula ChR2). OFC vs insula hotspot sites did not differ in levels of place-based self-stimulation at either 20Hz or 40 Hz frequencies (20 Hz Difference Score: *M = 153.3*, *SEM =* 62.4; 40 Hz Difference Score: *M = 236.8*, *SEM =* 41.3; No Laser Score: *M = -111.2*, *SEM =* 55.5; laser freq x brain site interaction: *F*2,73= 0.88, *p* =0.42), and male and female rats ChR2 rats with insula/OFC hotspot sites also showed comparable levels of self-stimulation, spending 3:2 to 2:1 duration ratios in laser-delivering chamber versus no-laser chamber, with no detectable sex difference (laser x brain site x sex interaction: *F*2,29 = 0.41 *p* = 0.67; *n* = 16 males *n* = 16 females). By comparison, eYFP control rats with OFC/insula hotspot sites failed to self-stimulate in the place-based task, spending equal amounts of time in both the non-laser and laser-delivering chambers (20 Hz Difference Score: *M = -119.28*, *SEM =* 91.3; 40 Hz Difference Score: *M = -102.2*, *SEM =* 97.9; No Laser Score: *M = -133.9*, *SEM =* 89.5; *n = 9* OFC eYFP, *n =* 4 insula eYFP),and so eYFP rats were different from ChR2 rats having similar hotspot sites (**Fig. 4, B**; **Fig. S9 A**; laser x virus interaction; *F*2,73= 3.69, *p* =0.03), *Coldstrip relatively fails to support place-based self-stimulation.* By contrast to OFC/insula hotspot place-based self-stimulation, and to coldstrip self-stimulation in the active-touch task, coldstrip ChR2 sites in the intervening caudal OFC and rostral insula, failed to reliably support place-based laser self-administration (**Fig 4, A, B**, **Fig S9, B**; laser main effect: *F*2,46= 0.46, *p* = 0.63). There did appear to be a non-significant trend toward place based self-stimulation at somes ites in caudolateral OFC (n=9), and opposite non-significant avoidance trends for some sites in in rostral to mid-insula (n=12), but these differences never reached p<.05 statistical significance as a group (Laser x site interaction; *F*2,46= 3.09, *p =* 0.06; caudolateral OFC ChR2 subgroup20 Hz Difference Score: *M = 58.8*, *SEM =* 57.4; 40 Hz Difference Score: *M = 246.7*, *SEM =* 39.2; No Laser Score: *M = -139.3*, *SEM =* 84.5; n= 9 anterior insula ChR2 subgroup 20 Hz Difference Score: *M = -204.0*, *SEM =* 49.8; 40 Hz Difference Score: *M = - 234.1*, *SEM =* 90.1; No Laser Score: *M = -35.92*, *SEM =* 61.53, *n* = 12 = insula ChR2). A potential reason for why results of the place based self-stimulation task might differ from active self-stimulation task for coldstrip Ch2 rats is that the cumulative duration of laser (seconds illumination per minute in laser-delivering chamber) was approximately 200% to 400% greater in the place-based task than in the active-touch task. It is possible that greater laser durations in the place-based task exceeded an optimal level for self-stimulation, and potentially became unattractive or even aversive at coldstrip sites in anterior insula and middle insula.

**Fig. 4.**
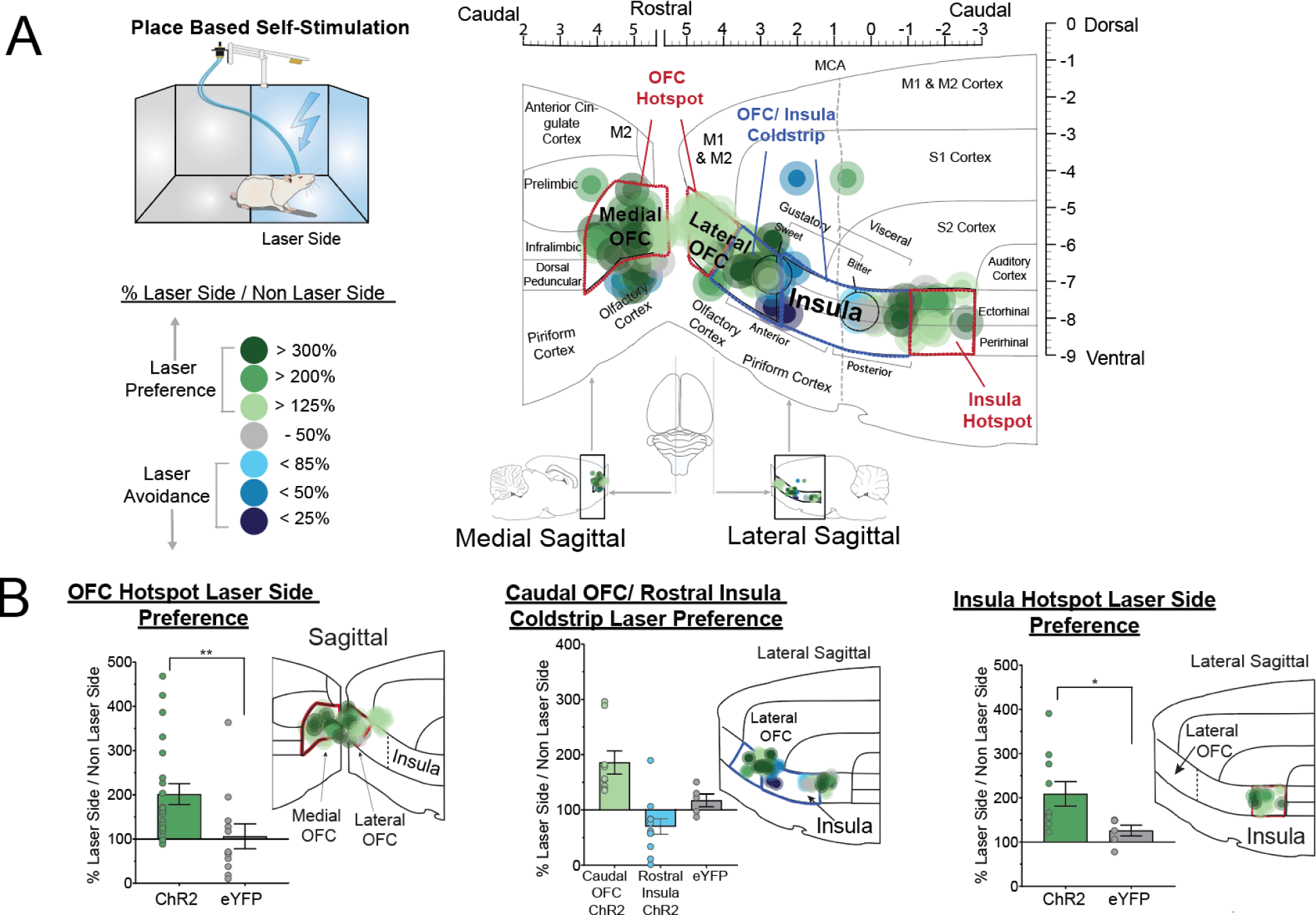
Cortical sites in orbitofrontal cortex and insula that support laser self-stimulation in place-based task. Caudal OFC and rostral to mid insula sites did not reliably promote place-based self-stimulation, and at some sites laser stimulation was avoided. (**A**) Functional maps show preferences for laser-paired side (green) or avoidance of laser-paired side (blue) during the place-based self-stimulation task. The color reflects the percent preference or avoidance for the laser-delivering vs. non-laser-side in the same rats. (**B**) Bar graphs show quantified % laser side preference in rostromedial OFC (left) and caudal insula (middle) ChR2 rats who showed evidence of self-stimulation relative to eYFP control rats (OFC Hotspot: *U*= 54.0, *Z* = -2.8, *p* = 0.004, *n* = 22 rostromedial OFC ChR2, *n* = 12 rostromedial OFC eYFP; Insula Hotspot: *U*= 10.0, *Z* = -2.2, *p* = 0.03, *n* =10 caudal insula ChR2; *n* = 6 caudal insula eYFP). No reliable place-based self-stimulation was observed in ChR2 rats with caudal OFC or rostral insula sites (*n* = 9 caudal OFC ChR2, *n* = 12 rostral OFC ChR2; *n* = 5 eYFP). All data presented as means and SEM, \**p*< 0.05; \*\**p*< 0.01.

Control eYFP virus rats failed to show either place based self-stimulation or laser avoidance in the place-based task(**Fig S9, B**; 20 Hz Difference Score: *M = 52.6*, *SEM =* 37.4; 40 Hz Difference Score: *M = 29.1*, *SEM =* 59.8; No Laser Score: *M = -11.38*, *SEM =* 120.6; laser x brain site x virus interaction: *F*2,46= 1.51, *p =* 0.23; *n = 3* OFC eYFP, *n =* 3 insula eYFP).

### Recruitment of Fos activation in distant limbic structures

We assessed distant changes in Fos protein expression in several mesocorticolimbic structures recruited by laser ChR2 excitation of neurons within cortical sites in OFC and insula. For all structures, distant Fos was measured after laser ChR2 illumination in a cortical site in either rostromedial OFC hotspot, far-caudal insula hotspot, or intervening coldstrip. Recruited Fos levels were compared with a) control eYFP Fos baseline levels measured in eYFP rats that received similar laser illuminations at corresponding cortical sites, and b) spontaneous baseline levels in control naïve rats that were lightly handled, but received no surgery, no virus or optic fiber, and no laser or behavioral testing. We particularly aimed to assess the hypothesis that neurobiological stimulation of neurons in a particular hedonic hotspot would recruit Fos elevation in distant hedonic hotspots in other limbic structures, producing unanimous activation in several hotspots simultaneously as a distributed hedonic network that may mediate ‘liking’ enhancements (*7*, *11*, *17*).

*Rostromedial OFC Hotspot neuron stimulation.* Laser ChR2 stimulation in the rostromedial OFC hotspot recruited distant 150%-300% increases in Fos expression in the caudal insula hotspot (**Fig. 5; Table S1)**. Similarly, OFC hotspot stimulation recruited distant ∼175%-300% increases in Fos in two other previously-identified subcortical hedonic hotspots, namely in NAc rostrodorsal medial shell, and in caudal ventral pallidum (**Fig. 5; Table S1)** (1–4, 7). By contrast, we did not observe OFC hotspot-induced increases in Fos in previously identified subcortical suppressive coldspots, such as the caudal subregion of the NAc medial shell or the anterior ventral pallidum. Significant Fos increases were observed also in prelimbic cortex, infralimbic cortex, anterior cingulate cortex, ventral tegmental area, nucleus accumbens core, perifornical areas of the lateral hypothalamus, and medial amygdala (**Fig. 5; Table S1)**.

**Fig. 5.**
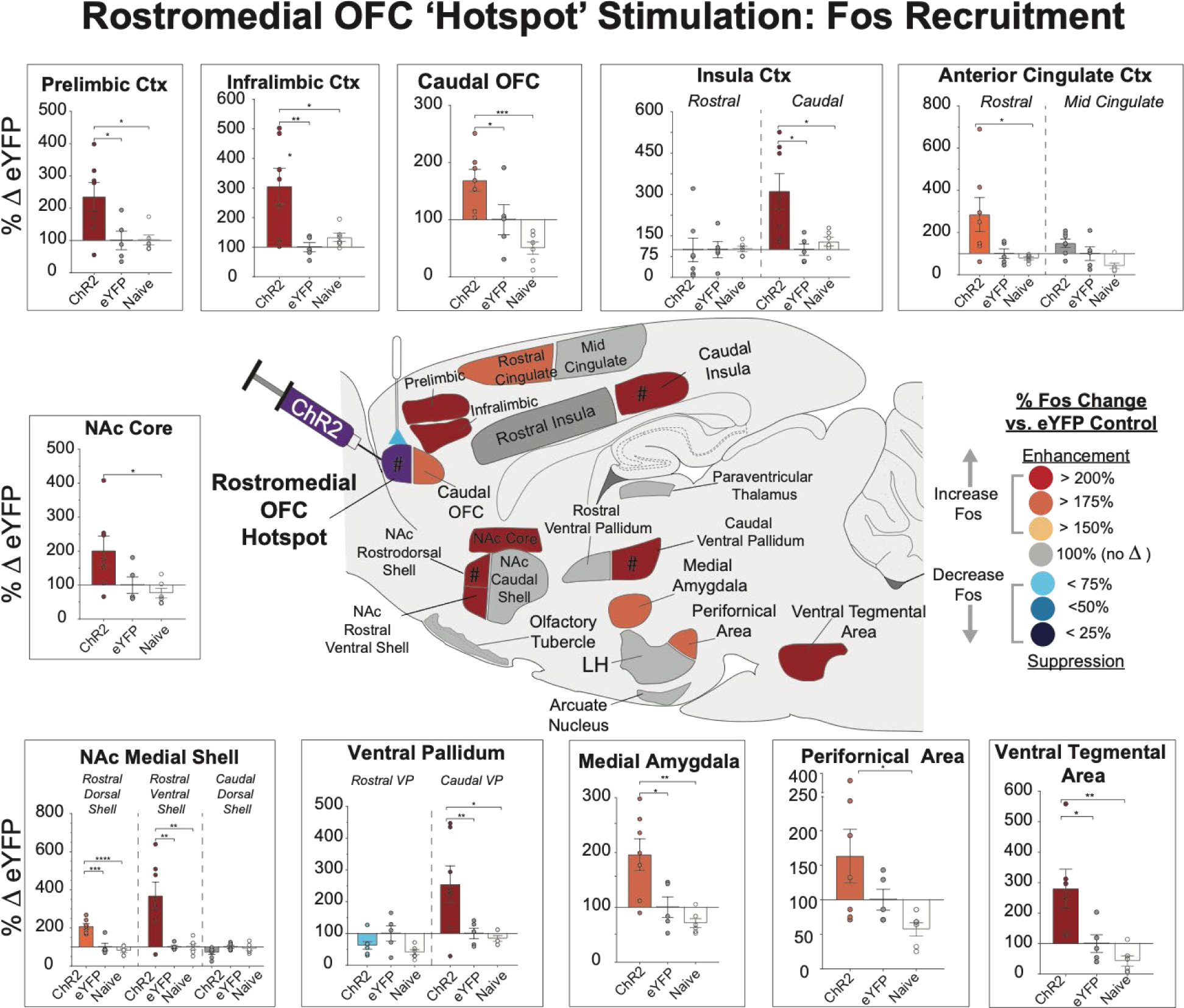
Distant Fos recruitment induced by rostromedial OFC ‘hotspot’. Optogenetic OFC hotspot stimulation recruits limbic brain circuitry for hedonic enhancement. Brain map shows elevated Fos expression in recruited mesocorticolimbic structures after laser stimulation in rostromedial OFC hotspot of ChR2 rats (*n* = 7; colors denote % Fos elevation compared to illuminated eYFP control rats (*n* = 5), and to naïve control baseline rats (*n* = 6). Significant Fos elevation was recruited in other hedonic hotspots, including far caudal insula cortex, nucleus accumbens rostrodorsal medial shell, and caudal ventral pallidum. Fos elevation was also recruited in other limbic cortical regions, such as prelimbic cortex, infralimbic cortex, caudal orbitofrontal cortex, and rostral anterior cingulate cortex. Fos elevation was also recruited in subcortical limbic structures, such as ventral tegmental area, nucleus accumbens core, nucleus accumbens rostroventral medial shell, medial amygdala, and perifornical area of the lateral hypothalamus. Also see Table S1. Bar graph data shown as mean and SEM of % Fos enhancements in that structure relative to eYFP controls. *p<0.05, **p<0.01, ***p<0.001, ****p<0.0001. # symbol denotes sites of previously identified hedonic hotspots (1, 2, 7).

*Caudal insula hotspot neuron stimulation.* Laser ChR2 stimulation in the far-caudal insula hotspot recruited distant >175% Fos increases in the rostromedial OFC hotspot **(Fig. 6, Table S2).** Insula hotspot stimulation also recruited >175% Fos increases in the subcortical hedonic hotspot in rostrodorsal quadrant of NAc medial shell, although no Fos change was detected in the posterior hotspot of ventral pallidum **(Fig. 6, Table S2)**. By contrast, Fos was not as elevated in suppressive coldspots of anterior ventral pallidum and rostral insula (though mild elevations were seen in caudal OFC and caudal NAc shell coldspots). Other significant increases in Fos were detected in prelimbic cortex, infralimbic cortex, anterior cingulate cortex, ventral tegmental area, nucleus accumbens core and shell, olfactory tubercle, paraventricular thalamus, lateral hypothalamus, and arcuate nucleus of hypothalamus **(Fig. 6, Table S2).**

**Fig. 6.**
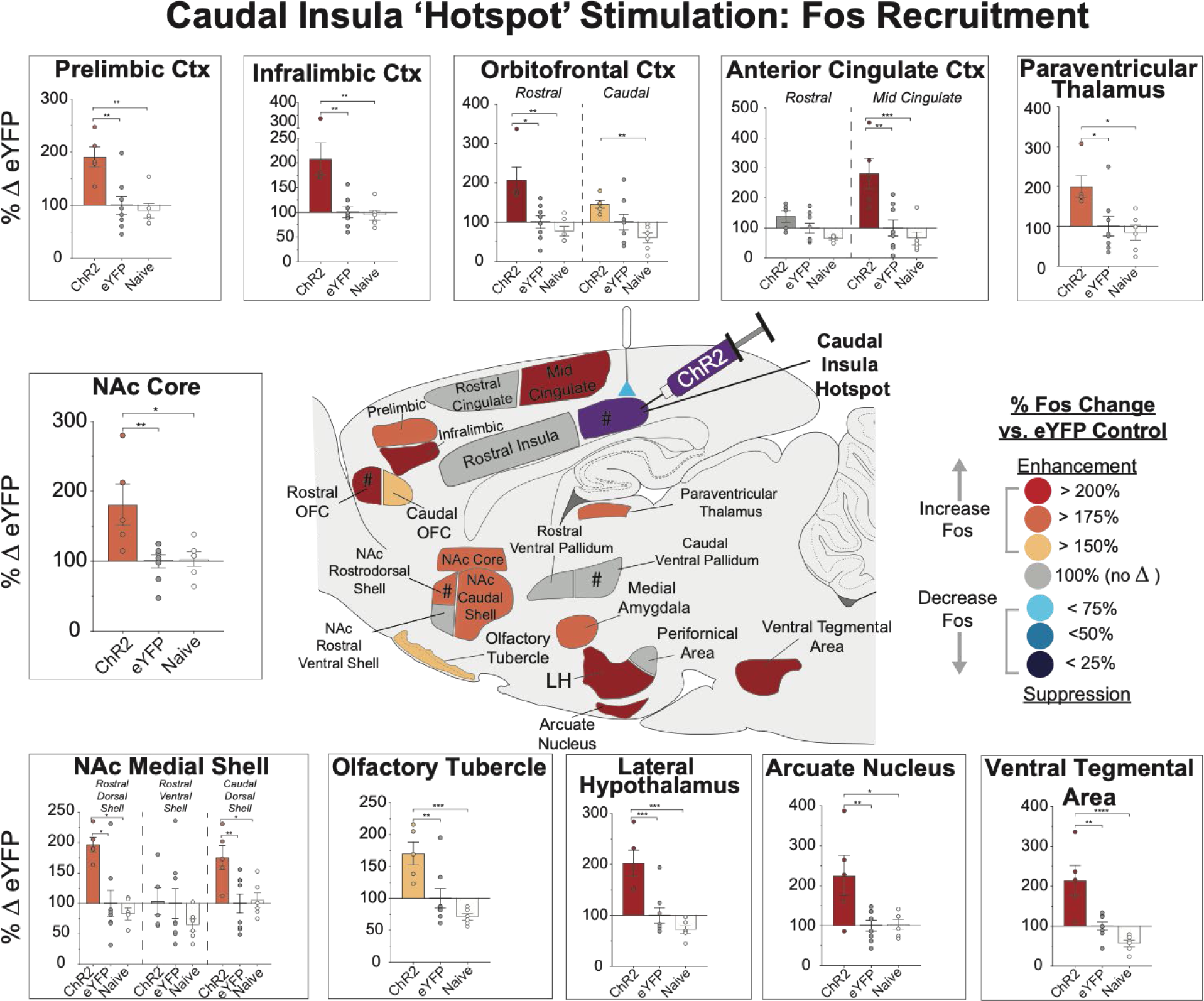
Distant Fos recruitment induced by caudal insula ‘hotspot’. Optogenetic stimulation in caudal Insula ‘Hotspot’ recruited limbic brain circuitry for hedonic enhancement. Brain map shows elevated Fos expression in recruited mesocorticolimbic structures after laser ChR2 stimulation in far-caudal insula hotspot (N = 5; colors denote % Fos elevation compared to eYFP control rats (N = 8), and to naïve control baseline rats (N = 6). Significant Fos elevation was recruited in other hedonic hotspots, including rostromedial orbitofrontal cortex, and nucleus accumbens rostrodorsal medial shell. Fos elevation was also recruited in other limbic cortical regions, such as prelimbic cortex, infralimbic cortex, caudolateral orbitofrontal cortex, and mid anterior cingulate cortex. Fos elevation was also recruited in other subcortical limbic structures, including ventral tegmental area, nucleus accumbens core, nucleus accumbens rostrocaudal medial shell, olfactory tubercle, paraventricular thalamus, lateral hypothalamus, and arcuate nucleus of ventromedial hypothalamus. Also see supplementary table 2. Bar graph data shown as mean and SEM of % Fos enhancements in that structure relative to eYFP controls. *p<0.05, **p<0.01, ***p<0.001, ****p<0.0001. # symbol denotes sites of previously identified hedonic hotspots (1, 2, 7).

Overall, our results support the idea that optogenetic neuronal stimulation in the rostromedial OFC hotspot is sufficient to also recruit Fos elevation in the far-posterior insula hotspot and vice versa. Laser ChR2 stimulation in the OFC hotspot or insula hotspot also simultaneously recruited significant Fos elevation in either one or both subcortical hedonic hotspots in NAc shell and ventral pallidum. This seems consistent with the hypothesis that local neurobiological stimulation of any one hedonic hotspot may recruit co-activation in other hotspots, to activate a more distributed hedonic enhancement circuit that enhances orofacial ‘liking’ reactions to sweetness.

*Caudal OFC to rostral-mid insula coldstrip.* By contrast, laser ChR2 stimulation in the suppressive hedonic coldstrip (i.e., sites in caudolateral OFC through anterior and middle insula) failed to recruit distant Fos increases in either cortical hedonic hotspot of rostromedial OFC or far-caudal insula. Similarly, cortical coldstrip stimulations failed to recruit Fos elevation in either subcortical hedonic hotspot in rostrodorsal medial shell or caudal ventral pallidum. However, stimulation in the hedonic coldstrip did increase Fos in other cortical coldstrip sites, as well as in other previously identified subcortical suppressive coldspots. For example, within the cortical coldstrip, caudolateral OFC stimulation recruited >200% Fos increases in rostral insula, and conversely rostral insula stimulations recruited >200% Fos increases in caudolateral OFC (**Fig. 7; Table S3**). Similarly, caudal OFC/rostral insula coldstrip stimulation also recruited >175% increases in the subcortical coldspot in NAc caudal medial shell, and recruited >200% Fos expression and recruited >200% Fos expression in the coldspot of rostral ventral pallidum (**Fig. 7; Table S3**) (all sites where opioid microinjections suppress hedonic ‘liking’ reactions to sucrose (1, 2, 4). Additional >175%-300% Fos increases were observed in prelimbic cortex, infralimbic cortex, ventral tegmental area, nucleus accumbens core, central amygdala, basolateral amygdala, hypothalamic arcuate nucleus, and paraventricular thalamus (**Fig. 7; Table S3**). Some of those additional activations may be related to the coldstrip-induced incentive motivational or ‘wanting’ effects found in the active-touch laser self-stimulation task.

**Fig. 7.**
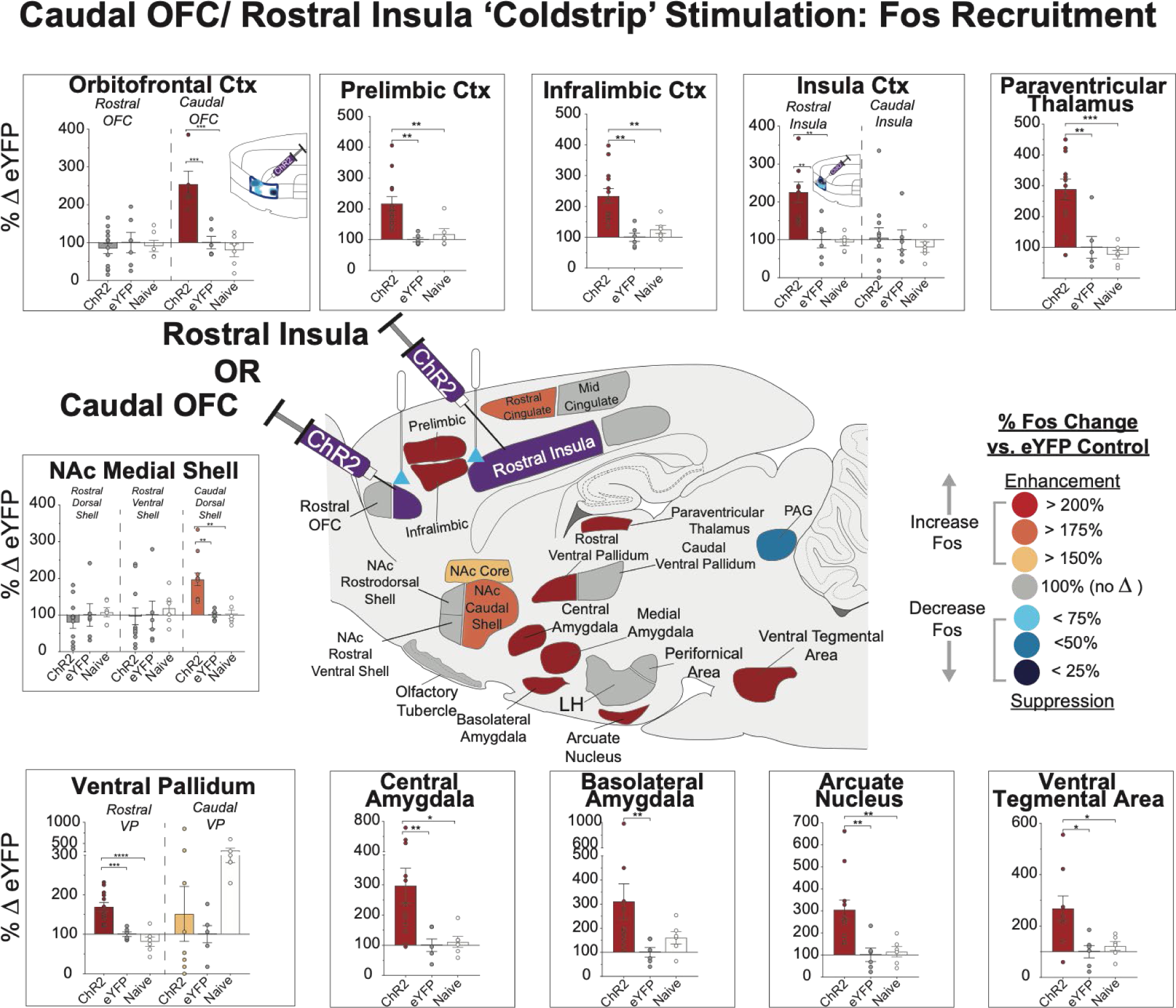
Caudal OFC/ rostral insula ‘coldstrip’ distant Fos recruitment. Brain map shows elevated Fos expression recruited in mesocorticolimbic structures after laser ChR2 stimulation in OFC/insula coldstrip ChR2 rats (N = 7 caudal OFC, N = 5 rostral insula; colors denote %Fos elevation compared to eYFP control rats (N = 6), and to naïve control baseline rats (N = 6). Cortical regions included caudal orbitofrontal cortex (Fos counts based on rostral insula ChR2 rats only), prelimbic cortex, infralimbic cortex, rostral insula cortex (Fos counts based on caudal OFC ChR2 rats only). Subcortical structures included nucleus accumbens core, caudodorsal nucleus accumbens medial shell, rostral VP, paraventricular thalamus, central amygdala, basolateral amygdala, arcuate nucleus, and ventral tegmental area. Also see supplementary table 3. Bar graph data shown as mean and SEM of % Fos Enhancements in that structure relative to eYFP controls. *p<0.05, **p<0.01, ***p<0.001, ****p<0.0001

## Discussion

Our results optogenetically confirm the existence, sites, and boundaries of two cortical hedonic hotspots able to causally enhance ‘liking’ reactions to sweetness, which were previously mapped only neurochemically (7). One optogenetic hedonic hotspot was confirmed in a rostromedial subregion of OFC and the other in a far-caudal subregion of insula. Our results extend understanding of their underlying neurobiological mechanisms by showing that optogenetic ChR2 stimulation within each approximately doubled the number of positive orofacial ‘liking’ expressions in rats elicited by sucrose taste, indicating an enhancement of hedonic impact. Our optogenetic results provide independent triangulating evidence that these hedonic hotspots are robust neurofunctional entities with special capacities to enhance the hedonic impact or ‘liking’ reaction to a pleasant stimulus, whether activated neurobiologically by optogenetic excitation here or by mu opioid agonist or orexin microinjections (7).

Anatomically, the locations and boundaries of the rostromedial OFC and caudal insula hotspots mapped optogenetically here were nearly identical to the boundaries previously mapped neurochemically(7), providing evidence that these anatomical subregions have special hedonic causal features. Our optogenetic mapping indicates the OFC hedonic hotspot begins anteriorly at the rostral tip of the OFC and extended caudally ∼2.5mm along both the medial and lateral surfaces of OFC. The insula hedonic hotspot was contained in the most posterior one-third of insula, approximately 5.3 mm^3^ in volume. In both hotspots, ChR2 laser excitation doubled the number of ‘liking’ expressions elicited by sucrose or water tastes. It may seem puzzling that the far-caudal subregion of insula containing its hedonic hotspot overlaps more closely with what is traditionally considered visceral sensory insula, rather than gustatory insula. However, a possible solution may lie in the phenomenon known as alliesthesia, in which relevant physiological states and their visceral signals modulate the pleasure of a sensory stimulus (18). For example, hunger vs satiety states modulate human subjective pleasure ratings and rat orofacial ‘liking’ reactions to sweetness. Potentially, visceral signals of hunger might be processed in the caudal insula hedonic hotspot to help mediate alliesthesia increase in sweetness ‘liking’.

In between the OFC hotspot and insula hotspot along the lateral surface of OFC, we also mapped a 4.5 mm long (17.0 mm^3^ volume) suppressive OFC-insula ‘hedonic coldstrip’. Throughout that coldstrip ChR2 laser excitations oppositely reduced the number of ‘liking’ reactions elicited by sweet tastes to about one-half control baseline levels (measured without laser in the same individuals). The suppressive coldstrip included the most-caudal one-third of OFC, and both the anterior insula and middle one-third of insula.

*Human parallels to hedonic hotspots?* In human neuroimaging studies, the rostromedial OFC hotspot in rats described here might correspond to a mid-anterior OFC site that is reported to track human subjective pleasantness ratings in fMRI studies, including changes in subjective taste pleasure induced by sensory-specific satiety (19–22). Similarly, human insula is reported to be activated by pleasant food images (23), and insula activation is reported to track decreases in chocolate taste pleasure ratings as people eat chocolate to satiety (24). A recent fMRI meta-analysis of human pleasure ratings for both beverages and humorous cartoons concluded “The spatial layout of the pleasure signature is consistent with… observations of hedonic hotspots identified in rodent studies”, and that human pleasure-coding fMRI sites also positively correlated with opioid binding (25). Other fMRI neuroimaging studies, as well as electrophysiological studies of nonhuman primates have implicated OFC and insula sites more generally in various aspects of reward (20, 21, 24, 26–29). Of course, hedonic hotspots were mapped here as causal sites in rats, namely as special subregions where local neuronal stimulations amplified ‘liking’ reactions to the hedonic impact of a pleasant stimulus, rather than as coding sites that activate spontaneously to the stimulus. Whether human pleasure-coding sites identified in fMRI studies also possess equivalent causal status as hedonic amplifiers remains an open question. Not all coding sites are necessarily also causal sites for hedonic impact (as some could take hedonic activation as an input to cause other nonhedonic functions). However, any apparent overlap between rat hedonic hotspots and human pleasure-coding sites may make those sites excellent candidates to amplify human pleasure.

*Motivational ‘wanting’ for laser anatomically more widespread than hedonic ‘liking’ enhancement.* By contrast to the restricted localization of hedonic hotspots for ‘liking’ enhancement, cortical sites that supported incentive motivational ‘wanting’ effects of ChR2 excitation extended between them into the hedonically-suppressive coldstrip, including posteriolateral OFC and anterior-mid insula sites, as well as OFC and insula hotspots, at least as measured by laser self-stimulation in the active-touch task. This is consistent with previous suggestions that cortical substrates for enhancement of motivational ‘wanting’ are anatomically more widespread than hedonic hotspots for ‘liking’, when ‘wanting’ rewards is measured as increases in food intake, instrumental responding during reward choice tasks, or self-administration of cortical electrical, or optogenetic, or drug microinjection stimulations (7, 13, 30–33).

*Comparison to other cortical optogenetic studies of hedonic taste modulation.* Previous studies reported that cortical optogenetic stimulation in anterior insula in mice enhanced voluntary licking of a drink spout and supported self-stimulation or elicited positive facial expressions to tastes (12–14). In partial agreement, we similarly found that anterior insula sites supported optogenetic self-stimulation in the spout-touch task. However, regarding hedonic impact, anterior insula sites here fell within our hedonic coldstrip where ChR2 excitation suppressed facial ‘liking’ reactions to sucrose. One possible explanation of the discrepancy is that most other studies primarily employed measures of reward motivation or ‘wanting’ rather than hedonic impact or ‘liking’.

The one exception was a study that used AI-scoring of positive facial reactions as mice licked taste solutions from a spout to conclude that anterior insula stimulation increased positive reactions (12). However, it is possible that the AI program included voluntary spout licking as a positive reaction, which could confound ‘liking and ‘wanting’ measures (the study did not report exactly which reactions were identified by the program). Voluntary licking is essentially a ‘wanting’ measure, as motivated licking is instrumentally required to ingest from an external source. Voluntary lick measures can give conclusions different from measures of ‘liking’ affective reactions to the hedonic impact of a taste delivered directly into the mouth, as measured here. For example, voluntary licking is reduced by s dopamine antagonists (34, 35), but dopamine suppression fails to reduce taste reactivity measures of hedonic ‘liking’ facial reactions to sweetness (36, 37), nor does dopamine suppression reduce human subjective liking ratings of sweet tastes or pleasant drugs even when it reduces wanting ratings of the same reward (38–40). If voluntary licking of an external sucrose source was included in an AI-coded positive score (12), then the anterior insula-induced increases could have reflected increased ‘wanting’ for sucrose, rather than increased ‘liking’.

Conversely, other optogenetic cortical-stimulation studies have reported that avoidance and aversive ‘disgust’ facial reactions in mice were evoked in caudal gustatory insula by optogenetic ChR2 stimulation (13, 14), including specifically stimulation of glutamate neurons using a CamKII promoter (41). However, the insula site that evoked ‘disgust’ in mice (13) might actually have been in the mid-insula portion of our hedonic coldstrip rather than in our far caudal insula hotspot, and in the mid-insula coldstrip we similarly promoted aversive ‘disgust’ reactions to sucrose. Further, that mouse study used laser intensities 10x higher than ours (10 – 20 mW vs 1-3 mW here)(13), and it’s conceivable that higher intensities in our insula coldstrip might have more robustly promoted aversive reactions. All those aversive effects contrast to our observation of enhancement of sucrose ‘liking’ reactions and laser self-stimulation for ChR2 excitation in the far caudal hotspot of insula in rats.

### Gains vs losses of hedonic function

In contrast to hotspot ChR2 enhancement of ‘liking’ reactions, iC++ inhibitions in the rostromedial OFC hotspot failed to reduce ‘liking’ reactions to sweetness or induce any other detectable change in taste-elicited reactions. Failure of hotspot inhibition to alter hedonic impact is consistent with reports that cortical lesions or even complete decortication in rats similarly fails to consistently impair taste ‘liking’ reactions or motivation for food reward (42, 43).

This difference between hotspot gain vs loss of hedonic function may reflect the hierarchical nature of cortical hedonic contributions to ‘liking’ reactions. That is, functional activation of cortical hedonic hotspot sites may cause hierarchical facilitation of subcortical hedonic circuits that generate positive ‘liking’ reactions to sweetness. Conversely, inhibition of cortical hotspots sites may remove hierarchical facilitation, but does not impair subcortical circuitry from semi-autonomously generating normal baseline ‘liking’ reactions. This lack of hedonic suppression from hotspot inhibition stands in contrast to suppression of ‘liking’ reactions by ChR2 stimulation of hedonic coldstrip sites. That suggests that coldstrip excitation hierarchically activates anti-hedonic subcortical circuitry of its own that actively suppresses or opposes ‘liking’ reactions.

### Potential Neuronal Mechanisms within Hedonic Hotspots

Our findings indicate that optogenetic depolarization of neurons within identified hotspot subregions of OFC and insula, induced by ChR2-mediated influx of Na+ and Ca+ ions, enhances hedonic ‘liking’ reactions to palatable tastes. This confirms anatomical identities for cortical hotspots previously reported by a study based on neurochemical opioid or orexin stimulations. However, that raises the mechanistic question of how both optogenetic excitation and microinjection of a mu opioid agonist or orexin in the same hedonic hotspot can cause similar hedonic enhancements? (7).

Orexin is reported to depolarize neurons in layer 6 of neocortex (44), as well as amygdala (45), nucleus basalis (46), and hypothalamus (47). Thus, orexin-induced neuronal depolarization in OFC or insula hotspots might conceivably underlie enhancement of ‘liking’, similarly to optogenetic ChR2 stimulation.

However, DAMGO is a selective mu-opioid agonist that acts at Gi-protein coupled inhibitory receptors to suppress intra-neuronal adenylyl cyclase and is associated with post-synaptic IPSPs (48–51). DAMGO-induced neuronal hyperpolarization would present a puzzle for understanding how both it and optogenetic depolarization in the same OFC or insula hedonic hotspots produce similar ‘liking’ enhancements. One possible resolution might be that mu-opioid microinjections inhibit local or afferent GABA inhibitory neurons, which disinhibits their target cortical neurons in the same site into depolarization, as has been proposed to occur in VTA, hippocampus, and periaqueductal gray (52–55). Consistent with this possibility, DAMGO is reported to reduce inhibitory synaptic transmission in insula cortex and ventrolateral and medial subregions of OFC (56, 57). An alternative possibility might be that a U-shaped polarization curve characterizes hedonic enhancement mechanisms within cortical hotspots. That would be similar to reports regarding the nucleus accumbens shell that both local inhibitory neuronal manipulations (e.g., GABA agonist microinjections; glutamate antagonist microinjections) (58–61) and local excitatory neuronal manipulations (e.g. optogenetic excitation; electrical stimulation) similarly increase appetitive motivation to eat (62–64). Clearly, future research is needed to solve this puzzle.

### Recruitment of Distant Hedonic Circuitry

Our measures of Fos elevation in distant limbic structures revealed that optogenetic stimulation in the rostromedial OFC hedonic hotspot recruited neurobiological Fos activation in neurons located in the other cortical insula hedonic hotspot, as well as in the subcortical hedonic hotspots of the NAc rostrodorsal medial shell and the caudolateral ventral pallidum (1, 2). Similarly, optogenetic stimulation of the far-caudal insula hedonic hotspot reciprocally recruited distant Fos activation in the rostromedial OFC hotspot, as well as also in the subcortical NAc rostrodorsal medial shell hotspot. The observation that stimulation of one hedonic hotspot recruits distant activation in multiple other hedonic hotspots is consistent with previous pharmacological studies (7, 11), and supports the hypothesis that ‘liking’ enhancement caused by stimulating any single hedonic hotspot may involve recruiting other hotspots into unanimous co-activation, as an entire integrated hedonic network. Although subcortical hedonic hotspots appear not to be directly connected anatomically (65, 66), both cortical and subcortical hedonic hotspots do appear to show functional connectivity or mutual co-activation, likely via intermediary sites, which may be crucial to enhancement of behavioral or psychological ‘liking’ reactions.

### Clinical Implications

Brain hedonic mechanisms that amplify ‘liking’ reactions to pleasant events may be relevant to understanding hedonic dysfunctions that may occur in depression and other affective disorders, which could aid efforts to improve clinical therapies for hedonic dysfunction (67–69). Some individuals with major depression or schizophrenia are reported to have symptoms of consummatory or true anhedonia, meaning inability to experience pleasure, while others may have a more selective avolition or anticipatory anhedonia, meaning loss of motivational ‘wanting’ for life rewards even if hedonic reactivity remains intact (70–73). A recent study in humans found that patients with major depressive disorder with anhedonia had blunted fMRI BOLD responses in OFC and insula during a monetary gain/ loss task, suggesting that in humans these cortical regions may contribute to hedonic dysfunction (74). Whether activation of the cortical suppressive ‘hedonic coldstrip’ described here contributes to reduced pleasure in true anhedonia, or whether promoting activity in hedonic hotspots could reverse such hedonic deficits remain open questions that could be addressed by future clinical research. Such possibilities would be in line with RDoC criteria, which breaks down psychological functioning into a subset of domains with underlying neurobiological determinants (75).

## Materials and Methods

### Animals

Female and male Sprague Dawley rats (*n =* 88; *n* = 44 female, *n =* 44 male; weighing 250-400 g at surgery), were group housed in separate same-sex rooms, maintained at 21° C constant temperature, on a reverse 12h dark/light cycle at the University of Michigan. *Ad libitum* access to both food and water was given throughout the experiments. Experimental procedures were approved by the Committee on the Use and Care of Animals at the University of Michigan.

### Surgery

*Optogenetic Virus Infusion.* Rats were anesthetized with isoflurane gas (4-5% induction, 1-2% maintenance) and received atropine (0.04 mg/kg; i.p.; Henry Schein) before surgery, and then placed into a stereotaxic apparatus (David Kopf Instruments). Bilateral microinjections either of AAV channelrhodopsin virus (ChR2: AAV5-hSyn-ChR2-eYFP; UNC Vector Core, Chapel Hill; 0.5 µL in insula sites - 0.75 µL in OFC sites) or of control virus lacking the opsin gene (eYFP: AAV5-hSyn-eYFP; UNC Vector Core, Chapel Hill, NC) were targeted at cortical sites in OFC (ChR2 *n* = 41; eYFP *n* = 13) and insula (ChR2 *n =* 19, eYFP *n* = 8) as described below. A separate group of rats received an inhibitory optogenetic virus (AAV5-iC++-eYFP; Stanford Vector Core; *n* = 7) in the rostromedial orbitofrontal cortex to determine whether neuronal inhibition in the OFC hedonic hotspot suppressed ‘liking’ reactions. At OFC sites a 0.75 µL volume of virus was infused per side over a 7.5-minute period at a constant rate of 0.1 µL/min. At insula sites, a lower 0.5 µL volume was infused per side, because pilot results indicated that 0.75 µL insula infusions may induce seizures during subsequent laser stimulation in ChR2 rats. Following virus infusion, the microinjector was subsequently left in place for an additional 10 min to allow for virus diffusion.

Sites were aimed to be as identical bilaterally as possible within each individual rat but were staggered across individuals so that the group’s sites filled the entire rostral-caudal extent of the lateral cortex from midline tip of anterior OFC to posterior insula. OFC coordinates ranged from +5.16 mm to + 3.00 mm AP, ± 0.2 to ±2.5 mm ML and -4.00 mm to -6.00 mm DV (all relative to bregma). OFC sites included medial orbitofrontal (MO) and ventral orbitofrontal (VO) subregions of medial OFC (± 0.2 to ±1.0 mm ML), and lateral to cover lateral orbitofrontal (LO), and dorsolateral (DLO) subregions of lateral OFC (± 1.5 to ±2.5 mm). Insula coordinates ranged from +3.00 to –1.56 mm AP, ± 4.00 to ±6.00 mm ML, and -5.00mm to -6.00 mm DV. Insula sites included anterior insula, middle insula and posterior insula subregions. After surgery, cefazolin (100 mg/kg, s.c.; Henry Schein) was administered to prevent infection, and carprofen (5 mg/kg, s.c.; Henry Schein) given for post-operative pain relief. Carprofen and cefazolin were repeated at 24-h and 48-h post-operation.

*Oral Cannula Surgery and Fiber Optic Implantation.* Three weeks after the initial viral infusion surgery, rats were re-anesthetized with isoflurane as described above for implantation of intracranial optic fibers and of bilateral oral cannulas, which allowed for direct oral infusions of sucrose, quinine, and water solutions. Each oral cannula (polyethylene-100 tubing) entered the upper cheek just lateral to the secondary maxillary molar, ascended beneath the zygomatic arch, and exited the skin at the dorsal head, where it was secured with skull screws and a dental acrylic headcap. In the same surgery, rats were implanted with bilateral optic fibers (200 µm), aimed to place each fiber tip 0.3 mm dorsal to the rat’s bilateral virus microinjection sites, and anchored with the same acrylic headcap. Cefazolin and carprofen were again administered and repeated post-operatively as above. All rats were allowed to recover for 1 week prior to behavioral testing.

### Stimulation Parameters and Order of Behavioral Tests

Laser stimulation was tested at 5 Hz, 10 Hz, 20 Hz, and 40 Hz frequencies (counterbalanced in order on a within-subject basis) at 1-3 mW intensity. Use of multiple frequencies assessed whether any effects of ChR2 stimulation were robust across a wide range, or instead limited to a particular frequency. Laser was always delivered bilaterally to OFC sites. Pilot insula results indicated that bilateral stimulation of insula sometimes produced seizures, and so unilateral laser stimulation was subsequently used at insula sites.

### Behavioral Procedures

*Taste Reactivity Testing.* Each rat was habituated to the test chamber for 30 minutes on four consecutive days before any behavioral testing occurred. On the last two days of habituation, rats received oral infusions of a 0.03M sucrose solution to habituate them to infusion of fluids into the mouth. In subsequent taste reactivity tests, affective orofacial reactions (i.e., positive ‘liking’ versus negative ‘disgust’ patterns) elicited by oral infusions either of water or of three different taste solutions: two concentrations of sucrose solutions (0.03M and 0.10M), and one concentration of bitter quinine (3 x 10^-4^ M) (8, 10). Orofacial reactions were videorecorded through a close-up lens facing an angled mirror underneath the transparent floor, positioned to capture a clear view of the mouth and face, and saved for subsequent offline analysis. Taste solutions (1 ml) were delivered into the mouth of rats through PE-50 tubing connected to a PE-10 delivery nozzle, at a constant 1ml/min rate during the 1 min infusion, via a syringe pump, connected to the oral cannula.

On each test day, a rat received two separate 1-ml/1-min infusions of the same solution (e.g., .01 M sucrose), one infusion accompanied by laser stimulation and the other infusion not accompanied by laser as a within-subject baseline (counterbalanced order across rats), spaced 8-10 min apart. Different tastants were tested on different days. During a laser-paired infusion, laser illumination (1-3 mW; 15 ms pulses) was cycled in 5-s ON, 5 Sec OFF bins throughout the 60-sec trial test. Several different frequencies of laser illumination within 5-s ON bins were tested on different days: Every laser parameter was tested on at least two days for each rat in separate daily tests.

*Taste Reactivity Scoring.* Taste reactivity videos were scored subsequently for positive hedonic ‘liking’ reactions, aversive ‘disgust’ reactions, and neutral taste reactions in slow-motion at speeds ranging from frame-by-frame to 1/5^th^ normal speed, using The Observer Software (Noldus; Leesburg, VA). Positive hedonic or ‘liking’ responses were considered to be: lateral tongue protrusions, paw licks, and rhythmic midline tongue protrusions (4, 7, 8). Aversive ‘disgust’ reactions were: gapes, forelimb flails, head shakes, face washes, chin rubs, and paw treading. Neutral responses (i.e., relatively uncoupled from hedonic impact) were: passive dripping of solution out of the mouth, rhythmic mouth movements, and grooming. A time-bin scoring system was used to ensure each type of affective reaction contributed equally to the overall affective score (4, 7, 8). Rhythmic mouth movements, paw licks, passive dripping, and grooming were all scored in 5-s time bins, because these behaviors typically are emitted in bouts of relatively long duration. Any continuous emission of these behaviors up to 5-sec was counted as a single occurrence; continuous emissions of 5-sec to 10-sec counted as two occurrences, etc. Midline tongue protrusions and paw-treading were scored similarly, but in 2-s bins, because they are typically emitted in shorter bouts. Lateral tongue protrusions, gapes, flails, headshakes, and chin rubs were counted as discrete events every time they occurred, because these can occur singly or in brief repetitions. A total positive hedonic (i.e., ‘liking’) score was then calculated by combining component scores of rhythmic tongue protrusions, paw licks, and lateral tongue protrusions. A total negative aversive (i.e., ‘disgust’) score was calculated by combining gapes, forelimb flails, head shakes, paw treading, face washes, and chin rubs (7, 8).

### Laser Self-Stimulation Tasks

To test whether laser ChR2 stimulation of cortical sites by itself would support incentive motivation for reward, in the absence of any taste stimulus, laser self-stimulation was assessed in two different tasks. A place-based self-stimulation task, similar to that used in early electrical brain-stimulation reward studies (76, 77), allowed rats to earn laser illuminations by entering a particular chamber in a 2-chamber apparatus and remaining there. Each side of the chamber was marked by a distinctive floor surface and different visual patterns on walls. Entry into the designated laser chamber triggered onset of laser stimulation (1-3 mW) at either 20 Hz or 40 Hz, depending on test day. Laser illumination continually cycled at 3-s ON, 8-s OFF as long as the rat remained within the designated laser chamber. Exit from the laser chamber terminated the laser pulses. Entry into the other chamber produced nothing.

Separately, an active-response or ‘spout-touch’ laser self-stimulation task allowed rats to earn brief laser illuminations each time they touched a particular one of two empty metal drinking spouts, positioned 5 cm apart on the wall of a Med-Associates operant chamber (Fairfax, VT). One spout was arbitrarily designated as the active ‘laser spout’, and each touch on it earned either a 1-s or 5-s duration bin (depending on day) of 15 ms laser pulses (1-3 mW) at either 20 Hz or 40 Hz (depending on day). Touches on a second inactive spout produced nothing and contacts on it simply served to measure baseline levels of exploratory touching. Spout assignments were balanced across rats. Each combination of laser parameters was repeated on 3 consecutive days of self-stimulation (30-min sessions, order of combinations balanced across rats).

### Immunohistochemistry and Histology

Beginning 75-min prior to euthanasia, a final laser stimulation session was administered with one of the laser parameters that produced hedonic modulation in the taste reactivity tests (40 Hz, 15 ms pulse, 5-sec ON/5-sec OFF; 30-min session). This final laser stimulation was given to a) induce local Fos plumes around optic fiber tips that would indicate the anatomical spread of local neuronal stimulation at that cortical site induced by ChR2 laser illumination, and b) potentially also recruit distant Fos activation in various limbic brain structures, to identify recruited circuitry that potentially might mediate optogenetic modulation of hedonic reactions (62).

Following the final laser stimulation, rats were deeply anesthetized with a lethal dose of sodium pentobarbital (150-200 mg/kg) and transcardially perfused with PBS followed by 4% PFA. Brains were removed and post-fixed in 4% PFA for 24-h and then transferred to a 25% sucrose solution for at least two days. Tissue was coronally sectioned at 40 micrometers using a cryostat (Leica), slices were processed for GFP and Fos protein immunohistochemistry, and imaged using a digital camera (Qimaging) and fluorescence microscope (Leica). For immunohistochemistry, coronal sections were rinsed for 10 min in 0.1 M sodium phosphate buffer three times, then blocked in 5% normal donkey serum / 0.2% triton-X PBS solution for 60 min and incubated overnight in a polyclonal rabbit anti-cfos igG primary antibody (1:2500; Synaptic Systems) and chicken polyclonal anti-GFP igY primary antibody (1:2000; Abcam). Tissue was again rinsed three times in 0.1M NaPb for 10 min followed by 2-h in biotin-SP-conjugated donkey anti-rabbit (1:300; Jackson ImmunoResearch) secondary antibody and AlexaFluor-488 donkey anti-chicken secondary antibody (1:300; Jackson ImmunoResearch). Tissue was rinsed three times in 0.1M NaPB for 10 min followed by1.5-h in streptavidin-conjugated Cy3 (1:300; Jackson ImmunoResearch). Brain sections were mounted, air-dried, and cover-slipped with anti-fade Pro-long gold (Invitrogen).

*Local Fos Plume Analysis.* Immunoreactivity for Fos-like protein was visualized using a fluorescent microscope filter with a band of excitation at 515-545 nm. Coronal sections were imaged (10x magnification) to localize fiber tips and surrounding Fos plumes, spread of virus expression, and to quantify Fos expression in distributed structures. Local Fos plumes, which are local Fos elevation induced by laser illumination that immediately surround an optic fiber tip, reflect how far local ChR2 neuronal excitation spreads (58). Fos plumes were mapped at 10x magnification by counting the number of Fos+ neurons within a 50 µm x 50 µm block sample of tissue, sampled consecutively along 8 radial arms emanating from the optic fiber tip. Counting continued outward along each arm until at least two consecutive blocks did not contain any Fos+ cells. This point determined the radius of the local Fos plume along that particular arm, and the same was done for all 8 arms. Percent increases in ChR2 Fos expression were calculated against a control baseline level measured at the same sites in eYFP rats with inactive virus control that also received laser illumination prior to euthanasia (to control for any Fos elevation merely due to local heat or light). Symbols matched to the size of observed Fos plumes were used to construct maps of ChR2 localization of function in OFC and insula figures (58, 62). Stimulation sites were plotted onto corresponding maps using a brain atlas (78).

*Recruitment of Fos changes in distant brain structures.* Functional activation of circuitry recruited by laser stimulation of OFC or insula immediately prior to euthanasia was assessed by measuring change in Fos expression at distant sites in multiple structures: infralimbic cortex, prelimbic cortex, orbitofrontal cortex, insula, anterior and posterior ventral pallidum, anterior and posterior nucleus accumbens shell and core, anterior and posterior lateral hypothalamus, anterior and posterior anterior cingulate cortex, central amygdala, basolateral amygdala, medial amygdala rostral and caudal ventral tegmentum, dorsolateral striatum, dorsomedial striatum, and paraventricular thalamus. For brain structures known to contain hedonic hotspots or coldspots (OFC, NAc, VP, and Insula) separate Fos counts were conducted in the hotspot and coldspot subregions of each structure. LASX software was used to capture tiled images of whole brain coronal sections at 10x magnification, using a filter with 515-545 excitation band. Within each subregion, Fos-expressing neurons were counted in two to three sample boxes, placed equidistantly within the structure, and approximately at the same locations across rats, guided by a template on a corresponding brain atlas to facilitate consistent box placements. The size of the sample boxes was adjusted to each brain structure, so that each box contained approximately 10 Fos+ neurons in naïve rats. Fos+ neurons were counted in each sample box by someone blind to experimental conditions. Fos counts across the 3 sample boxes were added together to determine expression for each subregion or structure.

### Statistical Analyses

Taste reactivity tests and spout self-stimulation tasks were analyzed using repeated-measures ANOVAs, following by *t-*tests for individual comparisons with a Bonferroni correction. Self-stimulation tasks were analyzed using mixed ANOVAs. Kruskal-Wallis tests and Friedman’s two-way ANOVAs were used for nonparametric tests, followed by Wilcoxon sign-ranked tests. Significance was set at *p* < 0.05.

## Acknowledgments

We thank Nina Mostovoi, Madeline Granillo, and Carina Castellanos for technical assistance, and Marc Bradshaw for equipment construction.

## Author contributions

Conceptualization: IM, KCB

Methodology: IM, KCB

Investigation: IM, KCB

Visualization: IM, KCB

Supervision: IM, KCB

Writing—original draft: IM, KCB

Writing—review & editing: IM, KCB

## Data and materials availability

All data are available in the main text or the supplementary materials. The data that support the findings of this study (Fig. 1-7) and supplement (Fig S1-S9; Tables S1-S3) will be made publicly available upon publication through NIH figshare public repository, https://doi.org/10.6084/m9.figshare.c.7356220.

Code for MED-PC software for spout self-stimulation tasks is also available through the NIH figshare public repository, https://doi.org/10.6084/m9.figshare.c.7356220.

