## Supplementary Materials for "Optogenetic hedonic hotspots in orbitofrontal cortex and insula: causing enhancement of sweetness ‘liking’"

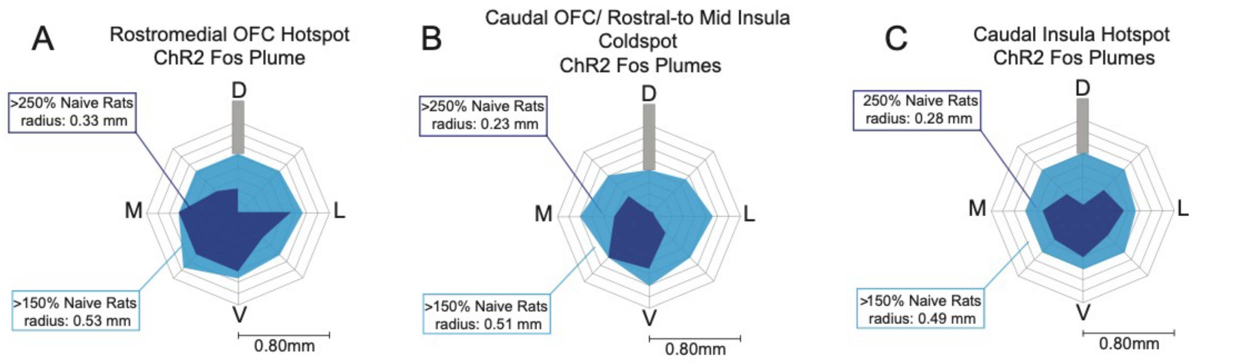


### Fig. S1.

**Cortical Hotspots Local Fos Plumes.** Local average Fos plumes around fiber tip measured after ChR2 laser stimulation at sites in (**A)** rostromedial OFC hotspot, **(B)** Caudal OFC and rostral to mid insula suppressive coldstrip, and (**C)** Far-caudal insula hedonic hotspot (Colors: >250% above naïve controls: light solid blue, > 150% above naïve control rats: dark solid blue).

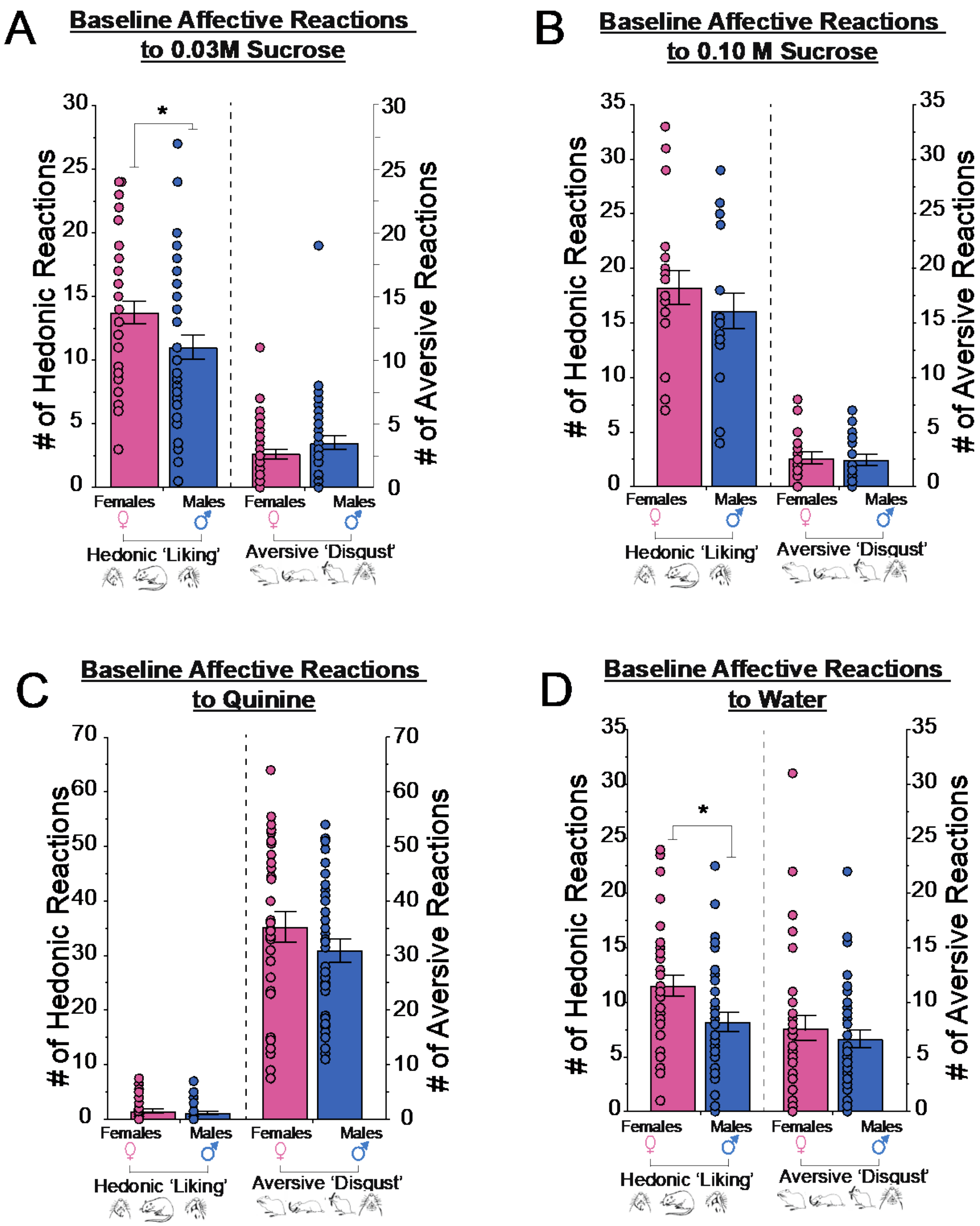


**Fig. S2. Sex Differences in taste reactivity at baseline.**

Sex differences and similarities in baseline affective taste reactivity elicited in male versus female rats in control condition without laser stimulation. **(A)** Females emitted higher positive 'liking' reactions to a dilute 0.03M sucrose solution ($n$ = 40 females, $n$ = 41 males). **(B)** No sex difference in positive hedonic reactions elicited by more concentrated 0.1M sucrose;(*n* = 20 females, *n* = 20 males). **(C)** No sex differences in negative ‘disgust’ reactions elicited by bitter quinine (*n* = 33 females, *n* = 35 males). **(D)** Females emitted higher positive ‘liking’ reactions to tap water infusions *(n* =33 females, *n* = 35 males). All data presented as mean and SEM. $*p < 0.05$.

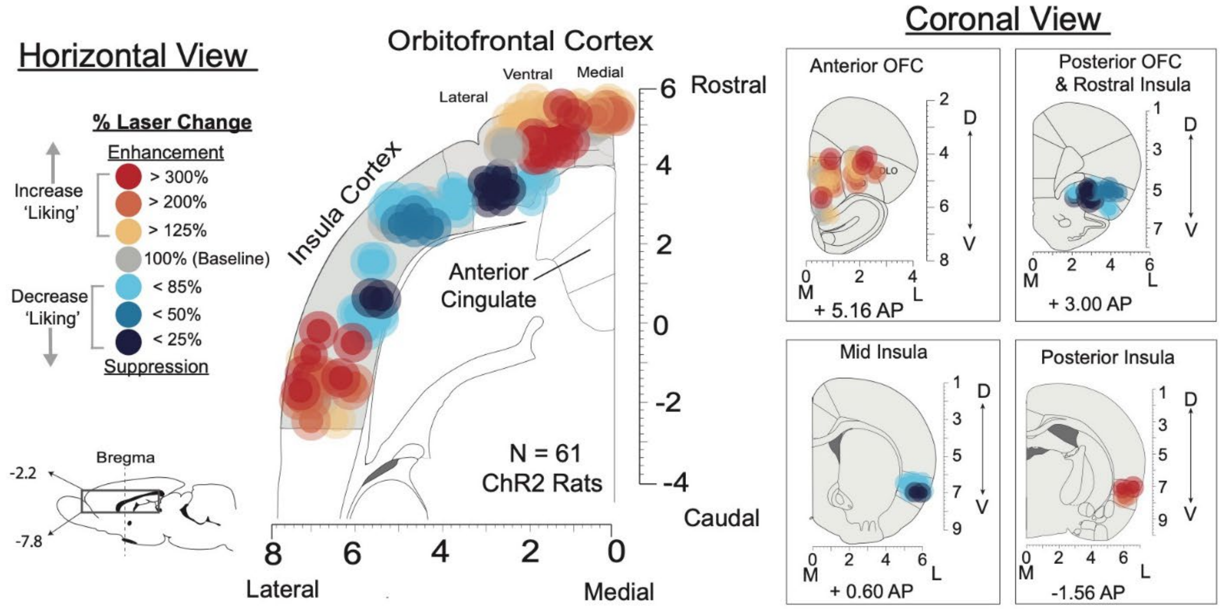


**Fig. S3. Coronal and Horizontal Localization of Function Maps**

Horizontal view (left) and coronal views (right) of hedonic hotspot localization of function. Each site shows laser ChR2 stimulation effects on positive 'liking' taste reactivity to sucrose. Each symbol placement reflects an individual rat. Size of symbols based on average size of Fos plumes. Color of each symbol represents the individual's within-subject change in hedonic reactions induced by ChR2 laser stimulation reflected as percent change from baseline hedonic reactions in no laser control condition in the same rats ('Liking' enhancements: red-yellow; 'Liking' suppression: Blue).

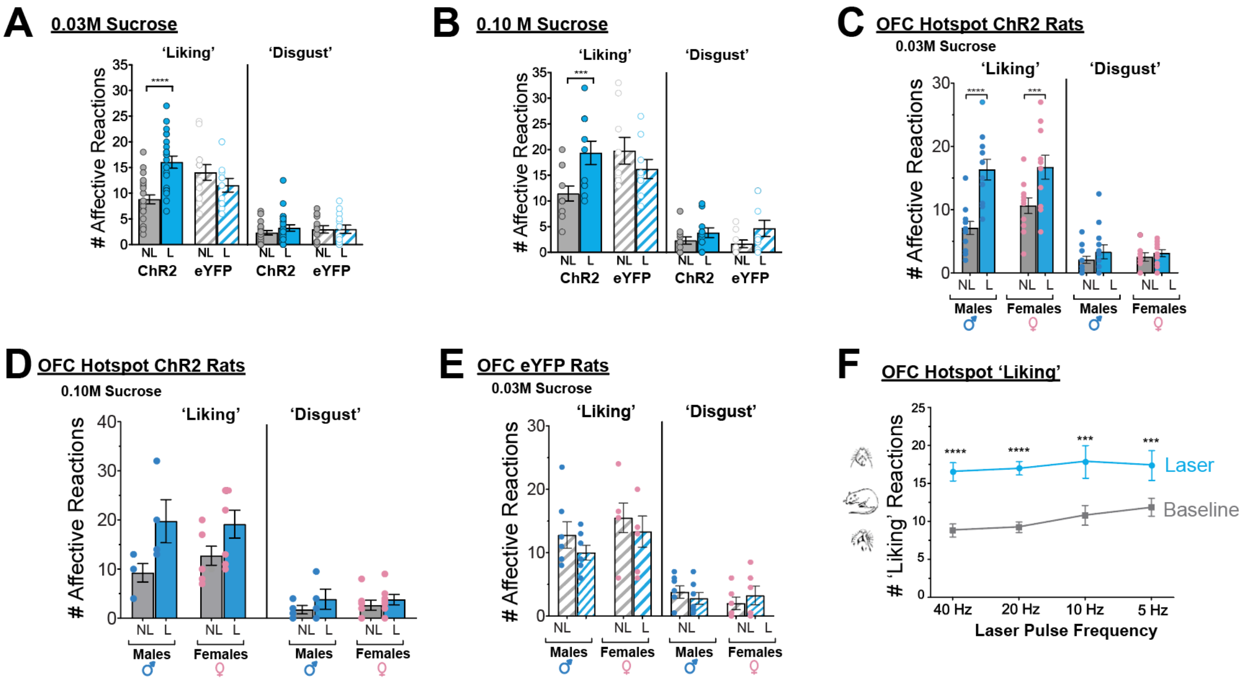


### Fig. S4. Rostromedial OFC Hotspot Sucrose Taste Reactivity

Raw affective components counts elicited by various sucrose concentrations following rostromedial OFC hotspot neuron activations. (**A**) Optogenetic laser activations increase hedonic 'liking' reactions to 0.03M sucrose in ChR2 rats ($n$ = 23), but not eYFP controls ($n$ = 23). (**B**) Laser activations increase hedonic 'liking' reactions to 0.1M sucrose in ChR2 rats ($n$ = 11), but not eYFP controls ($n$ = 23). (**C**) Male ($n$ = 12) and female ($n$ = 11) rostromedial OFC hotspot ChR2 rats show similar laser-induced increases in positive 'liking' reactions to 0.03M sucrose. (**D**) Similar positive 'liking' reaction enhancements in OFC hotspot ChR2 male ($n$ = 4) and female rats ($n$ = 7) to 0.10M sucrose. (**E**) No sex differences in affective expressions to 0.03M sucrose in eYFP control rats ($n$ = 13). (**F**) Multiple laser frequencies (5, 10, 20, 40 Hz; all 1 mW) increase positive 'liking' reactions to 0.03M sucrose in ChR2 rats. Data shown as within-subjects percent change from no laser baseline conditions. 40 Hz: U= 17.0, Z= -4.4, p < 0.001; 20 Hz; U= 31.5, Z= -3.44, p < 0.001; 10 Hz: U= 18.0, Z= -2.59, p = 0.008; 5 Hz: U= 20.5, Z= -2.42, p = 0.015, All data shown as means ± SEM. *$p$< 0.05, ***$p$<0.001.

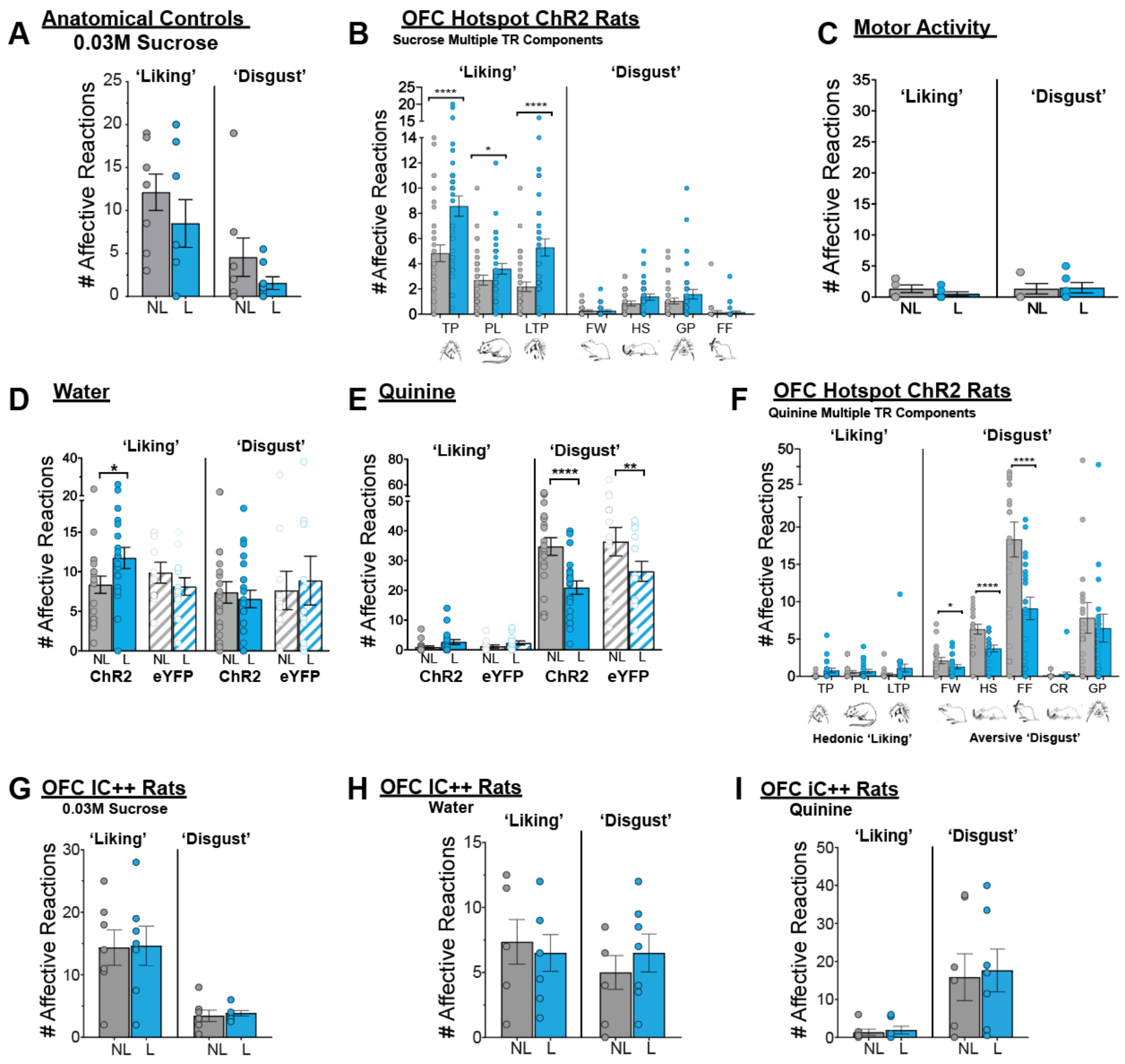


### Fig. S5. OFC Hotspot Taste Reactivity

Raw affective components counts elicited by various sucrose concentrations following rostromedial OFC hotspot neuron activations. (**A**) Laser ChR2 activations in anatomical control rats (prelimbic cortex and olfactory cortex; $n = 7$) fail to increase affective reactions to 0.03M sucrose. (**B**) Positive hedonic taste reactivity components include: paw licks (PL), lateral tongue protrusions (LTP), and rhythmic tongue protrusions (TP). Negative 'disgust' components are: gapes (GP), face washes (FW), head shakes (HS), forelimb flails (FF), and chin rubs (CR). Relative neutral components are: rhythmic mouth movements and passive dripping (not shown). Scoring: each occurrence was counted for LTP, GP, HS, FF, and CR. TP was scored in 2-s bins, and PL was scored in 5-s bins. Bar graphs show absolute scores as mean and SEM for no laser baseline and ChR2 laser stimulation trials in the same ChR2 rats (grey bars: no laser trials; red and blue bars: laser trials). Laser ChR2 stimulation (40 Hz) in rostromedial OFC hotspot sites significantly increased TP, PL, and LTP hedonic reactions to 0.03M and 0.10M sucrose collapsed together (laser x concentration interactions: TP: $F_{1,32} = 0.48$, $p = 0.49$; PL: $F_{1,32} = 0.09$, $p = 0.80$; PL: $F_{1,32} = 0.51$, $p = 0.48$. (**C**) No oromotor reactions observed from OFC hotspot ChR2 ($n = 6$) laser activations in the absence of taste infusions (**D**) Rostromedial OFC ChR2 ($n = 21$) activations increase hedonic 'liking 'reactions to tap water and not in eYFP controls ($n = 12$). (**E**) Aversive 'disgust' reactions to quinine are reduced by rostromedial OFC laser stimulation in both rostromedial ChR2 ($n = 21$) and eYFP rats ($n = 13$). (**F**) Laser ChR2 stimulation in rostromedial

OFC hotspot decreased aversive 'disgust' reactions of FW, HS, and FF elicited by bitter quinine. (**G-I)** Rostromedial OFC hotspot optogenetic iC++ inhibitions fail to alter affective reactions elicited by 0.03M sucrose, water, or quinine. All data shown as means ± SEM. *$p<0.05$, **$p<0.01$, ***$p<0.001$, ****$p< 0.0001$.

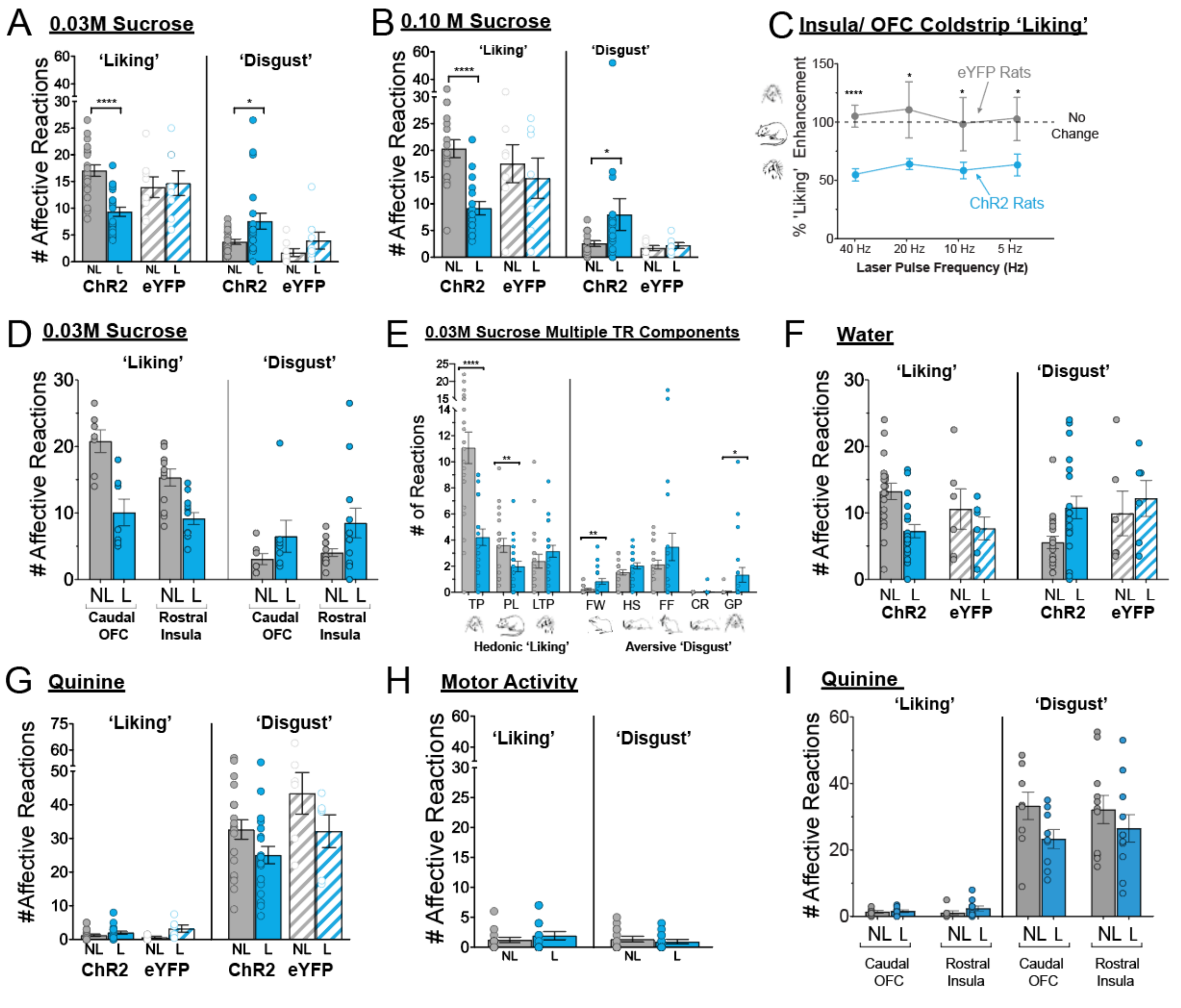


**Fig. S6. OFC/ Insula Coldstrip Taste Reactivity.**

Raw affective component counts elicited by various tastants following optogenetic activations of neurons in the caudal OFC/ rostral-to-mid insula hedonic 'coldstrip'. (**A**) Optogenetic laser activations decrease hedonic 'liking' reactions and increase aversive 'disgust' reactions to 0.03M sucrose in ChR2 rats (*n* = 17), but not in eYFP controls (*n* = 6) (**B**) Laser ChR2 activations suppress hedonic 'liking' reactions to 0.1M sucrose (**C**) Multiple laser frequencies (5, 10, 20, 40 Hz; all 1 mW) suppress positive 'liking' reactions to 0.03M sucrose in ChR2 rats. Data shown as within-subjects percent change from no laser conditions. (**D**) Laser ChR2 activations similarly suppress hedonic 'liking' reactions to 0.03M sucrose in OFC and Insula ChR2 rats. (**E**) Positive hedonic reactivity components include: paw licks (PL), lateral tongue protrusions (LTP), and rhythmic tongue protrusions (TP). Negative 'disgust' components are: gapes (GP), face washes (FW), head shakes (HS), forelimb flails (FF), and chin rubs (CR). Relative neutral components are: rhythmic mouth movements and passive dripping (not shown). Scoring: each occurrence was counted for LTP, GP, HS, FF, and CR. TP was scored in 2-s bins, and PL was scored in 5-s bins. Bar graphs show absolute scores as mean and SEM for no laser baseline and ChR2 laser stimulation trials in the same ChR2 rats (grey bars: no laser trials; blue bars: laser trials). Laser ChR2 stimulation in caudal OFC - rostral insula coldstrip sites oppositely suppressed TP and PL hedonic 'liking' reactions to sucrose and increased aversive 'disgust' FW and GPs. (**F)** Laser ChR2 activations in caudal OFC and rostral insula coldstrip sites decrease positive 'liking' reactions to water in ChR2 9 (*n* = 20) and eYFP controls (*n* = 6). (**G**) Laser activations in caudal OFC and rostral insula coldstrip sites decrease aversive 'disgust' reactions to quinine in ChR2 (*n* = 20) rats and eYFP controls (*n* = 6). (**H**) No oromotor reactions observed from caudal OFC and rostral insula coldstrip activations in the absence of taste in ChR2 rats (*n* = 14). (**I**) Laser ChR2 activations in either caudal OFC segment or rostral insula segment of intervening coldstrip similarly suppress aversive 'disgust' reactions to quinine. All data shown as means ± SEM. $*p<0.05$, $**p<0.01$, $***p<0.001$, $****p<0.0001$

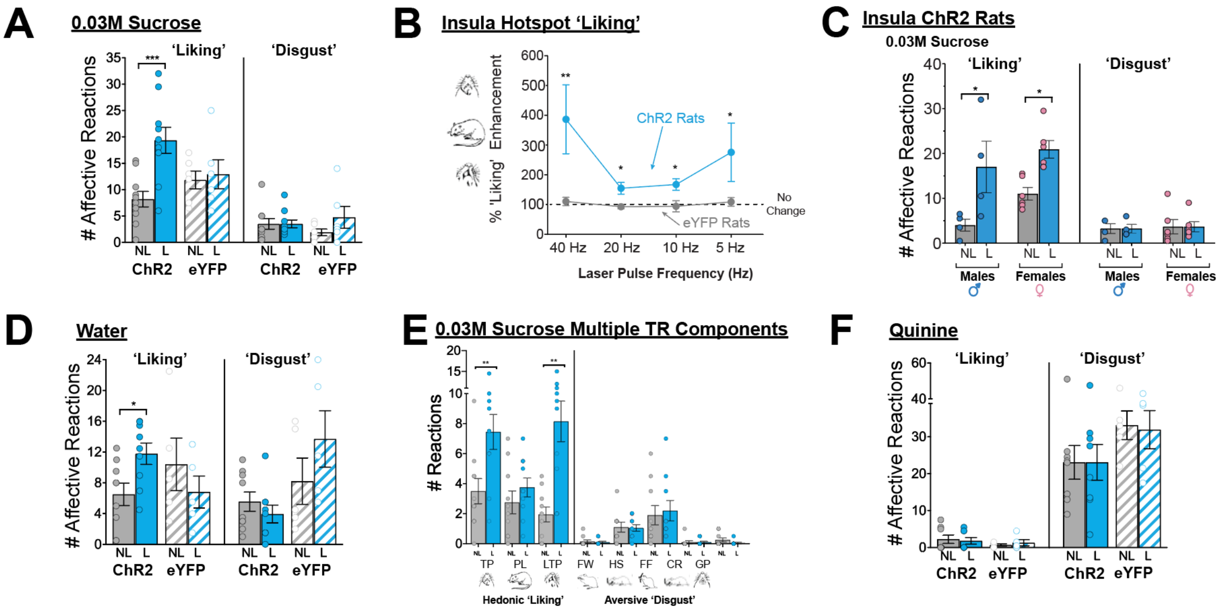


**Fig. S7. Insula Hotspot Taste Reactivity**

Raw affective component counts elicited by various tastants following optogenetic activations in the far-caudal insula hedonic 'hotspot'. (**A**) Optogenetic laser activations increase hedonic 'liking' reactions to 0.03M sucrose in ChR2 rats ($n$ = 10), but not eYFP controls ($n$ = 6). (**B**) All laser frequencies (40 Hz, 20 Hz, 10 Hz, and 5Hz; all 1 mW) increased positive 'liking' reactions to 0.03M sucrose in insula ChR2 rats. All data shown as laser induced percent increases from no laser conditions in the same animals; 40 Hz: U= 4..0, Z= -2.82, p = 0.003; 20 Hz; U= 0.00, Z= -3.1, p = 0.001; 10 Hz: U= 5.30, Z= -2.1, p = 0.05; 5 Hz: U= 6.0, Z= -2.1, p = 0.04. (**C**) Male ($n$ = 4) and female ($n$ = 6) rats with sites in caudal insula hotspot show equal laser-induced ChR2 increases in hedonic reactions to sucrose. (**D**) Caudal insula hotspot ChR2 laser activations increase positive 'liking' reactions to water. (**E**) Laser ChR2 stimulation in far-caudal insula hotspot increased both TP and LTP positive 'liking' reactions elicited by 0.03M sucrose. (**F**) No change in aversive 'disgust' reactions to quinine following caudal insula ChR2 activations. All data shown as means ± SEM. *$p$<0.05, **$p$<0.01, ***$p$<0.001

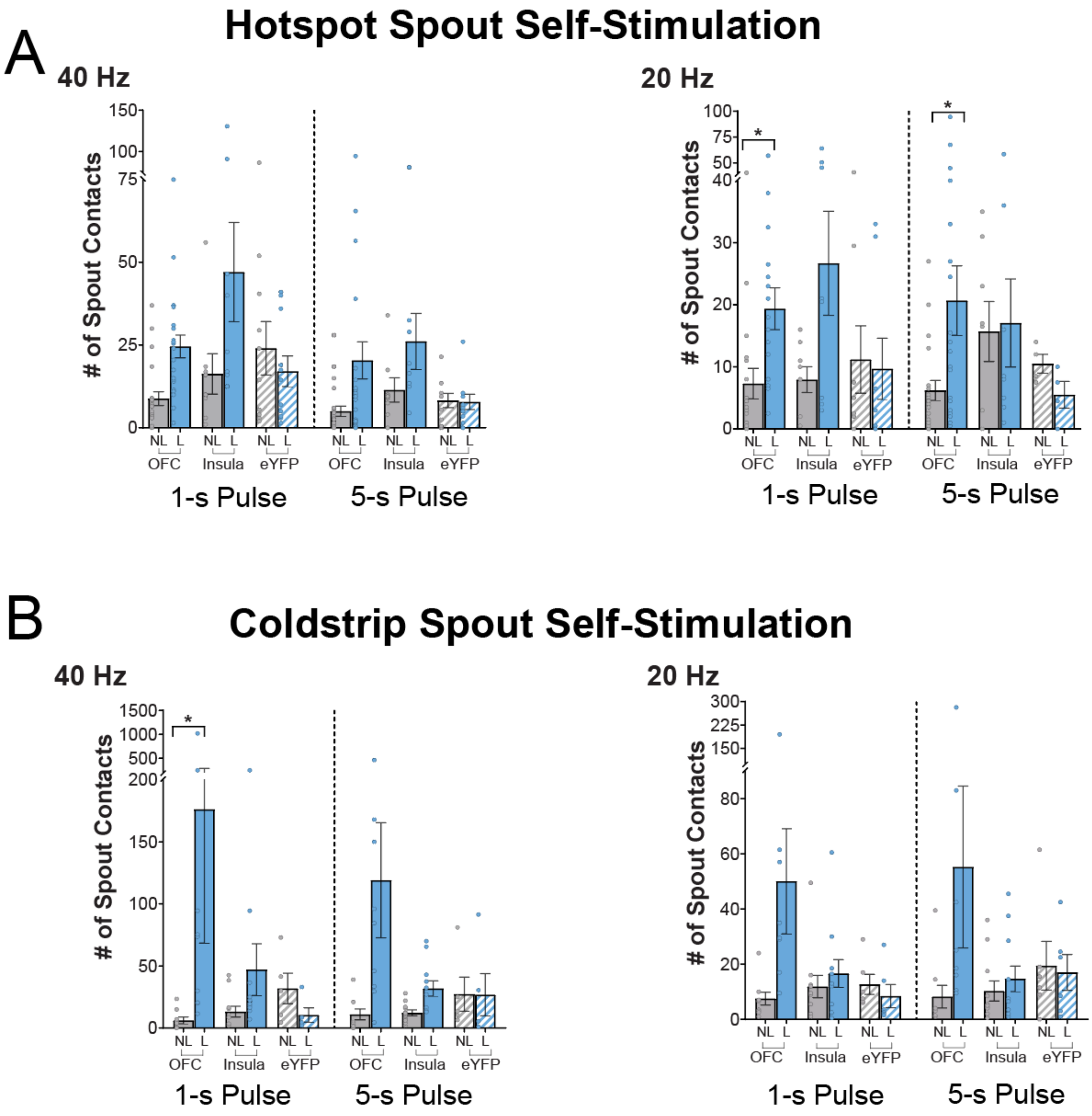


**Fig. S8. OFC and insula spout self-stimulation.**

**(A)** Total laser self-stimulations earned on spout-touch task by ChR2 rats with rostromedial and caudal insula hedonic hotspot sites (combined) at both 40 Hz (left) and 20 Hz (right) laser frequencies (5 sec and 1 sec pulse durations; 1 mW). (**B**) Total laser self-stimulations earned by rats from caudal OFC and rostral-to-mid insula coldstrip sites at both 40 Hz (left) and 20 Hz (right) laser frequencies (5 sec and 1 sec pulse durations; 1 mW). All data presented as means and SEM; *$p < 0.05$.

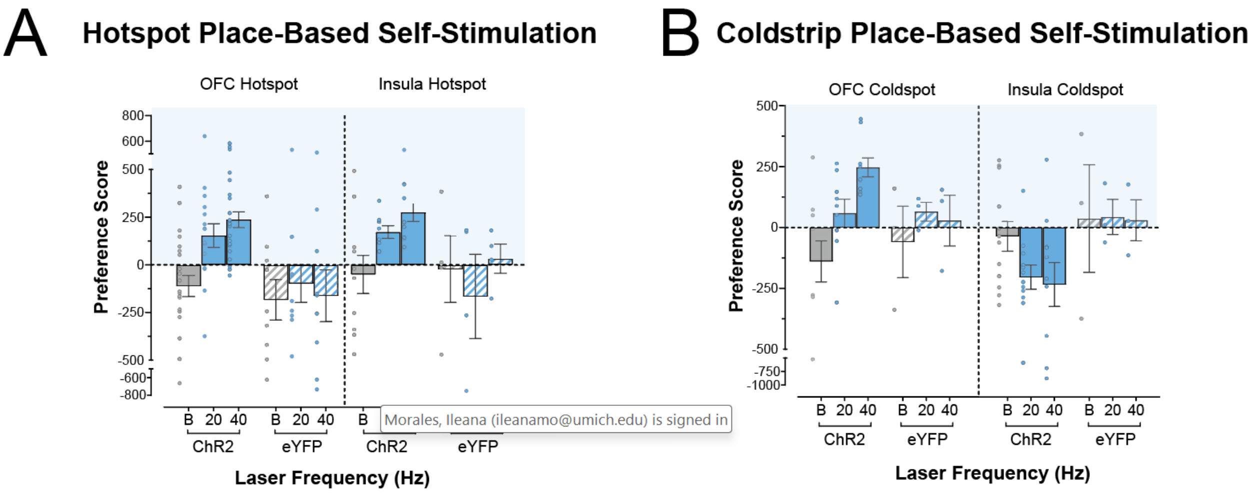


**Fig. S9.**

**OFC and insula place-based self-stimulation. (A)** Raw preference scores during place-based self-stimulation tests for ChR2 rats with sites in rostromedial and caudal insula hedonic hotspots (combined) (*n* = 22 rostromedial OFC ChR2; *n* = 9 rostromedial OFC eYFP; *n* =10 caudal insula ChR2; *n* = 4 caudal insula eYFP). **(B)** Raw preference scores during place-based self-stimulation tests for for ChR2 rats with sites in caudal OFC and rostral to mid insula coldstrip (*n* = 9 caudal OFC ChR2; *n* = 3 caudal OFC eYFP; *n* = 12 rostral to mid insula ChR2; *n* = 3 rostral insula eYFP). All data presented as means and SEM; Preference score reflects time (s) spent on the laser side – time (s) spent on non-laser side in the same rats; B: baseline habituation day; 20: 20 Hz stimulation tests; 40: 40 Hz stimulation tests.

| OFC Hotspot Fos + Neurons | Fos+ Counts (MEAN ±SEM) | | | Statistic | | Bonferroni Adjusted vs eYFP | 95% CI | Effect Size |
|---|---|---|---|---|---|---|---|---|
| | ChR2 | eYFP | Naïve | | | | | |
| Brain Region | n = 7 | n = 5 | n = 6 | *F (ANOVA) or H*$^{\#}$ *(Kruskal- Wallis)* | *p* | *p* | | *d or* $\eta^{2\#}$ |
| Prelimbic Cortex | 136.9 ± 25.9 | 58.2 ± 16.8 | 59.0 ±8.9 | 5.43 | 0.017* | 0.02* | (17.4, 139.9) | 1.4 |
| Infralimbic Cortex | 122.4 ± 24.2 | 40.0 ± 6.2 | 53.3 ± 5.6 | 7.03 | 0.001** | 0.004** | (30.2, 134.7) | 1.78 |
| Posterior OFC | 143.1 ± 16.4 | 84.8 ± 22.4 | 42.7 ± 9.1 | 10.43 | 0.001** | 0.02* | (8.6, 108.1) | 1.2 |
| Anterior Insula | 45.0 ± 19.3 | 45.4 ±13.2 | 46.5 ± 4.5 | 1.12# | 0.59 | | | |
| Posterior Insula | 122.1 ± 25.0 | 39.2 ± 8.1 | 50.7 ± 6.4 | 6.64 | 0.009* | 0.006** | (28.2, 137.7) | 1.71 |
| Rostral Anterior Cingulate | 111.3 ± 31.4 | 39.0 ± 8.9 | 30.3 ± 6.7 | 5.86# | 0.04* | 0.06 | | |
| Mid Anterior Cingulate | 59.7 ± 7.8 | 40.0 ± 13.0 | 16.8 ± 5.4 | 6.58 | 0.009** | 0.13 | | |
| NAc Core | 70.1 ± 15.0 | 34.8 ± 8.3 | 26.5 ± 4.9 | 4.6 | 0.03* | 0.04* | (0.75, 69.9) | 1.1 |
| NAc Dorsal Medial Shell | 106.9 ± 7.1 | 51.2 ± 9.8 | 41.5 ± 4.8 | 25.5 | <0.0001*** | <0.0001**** | (33.5, 77.85) | 2.7 |
| NAc Ventral Medial Shell | 84.7 ± 16.7 | 23.0 ± 1.8 | 23.3 ± 3.6 | 9.96 | 0.002** | 0.002** | (26.1,97.3) | 2.0 |
| NAc Caudal Dorsal Medial Shell | 28.4 ± 4.0 | 40.0 ± 3.4 | 36.7 ± 4.1 | 2.3 | 0.13 | | | |
| Dorsolateral Striatum | 24.6 ± 11.8 | 22.0 ± 5.0 | 12.7 ± 1.6 | 0.57 | 0.58 | | | |
| Olfactory Tubercle | 32.4 ± 10.5 | 20.2 ± 7.3 | 13.8 ± 0.9 | 1.5 | 0.25 | | | |
| Anterior VP Hotspot | 26.1 ± 5.0 | 41.8 ± 10.1 | 17.2 ± 3.4 | 3.7 | 0.05 | | | |
| Posterior VP Hotspot | 46.1 ± 8.3 | 17.0 ± 2.8 | 14.3 ± 1.5 | 9.6 | 0.002** | 0.004** | (11.1, 47.22) | 2.0 |
| Bed Nucleus Stria Terminalis | 20.7 ± 3.6 | 27.4 ± 7.9 | 26.5 ± 3.0 | 0.61 | 0.56 | | | |
| Lateral Hypothalamus | 36.7 ± 9.4 | 29.0 ± 6.9 | 20.8 ± 2.1 | 3.32# | 0.2 | | | |
| Perifornical Area | 45.7 ± 10.8 | 28.0 ± 4.2 | 16.0 ± 2.7 | 3.98 | 0.04* | | | |
| Central Amygdala | 40.1 ± 15.8 | 20.6 ± 4.9 | 15.2 ± 2.5 | 2.30# | 0.33 | | | |
| Basolateral Amygdala | 32.0 ±5.7 | 29.0 ±8.5 | 27.8 ± 4.5 | 0.13 | 0.87 | | | |
| Medial Amygdala | 63.1 ± 9.3 | 32.2± 6.1 | 23.0 ± 2.6 | 3.48 | 0.002** | 0.008** | (9.2,52.7) | 1.5 |
| Paraventricular Thalamus | 23.7 ± 2.8 | 17.2 ± 4.0 | 16.17 ± 3.0 | 1.76 | 0.21 | | | |
| Arcuate Nucleus | 21.0 ± 6.0 | 15.4 ± 2.1 | 20.2 ± 4.1 | 0.35 | 0.71 | | | |
| Substantia Nigra | 20.4± 6.1 | 17.6 ± 2.4 | 7.3 ± 3.2 | 5.56# | 0.06 | | | |
| Ventral Tegmental Area | 58.6 ± 11.2 | 20.2 ± 5.9 | 8.7 ± 3.4 | 12.24# | 0.0001*** | 0.04* | | 0.68# |
| Periacqueductal Gray | 18.2 ± 3.7 | 18.6 ± 2.5 | 11.8 ± 0.8 | 2.1 | 0.16 | | | |

### Table S1.

**Raw Fos counts after rostromedial OFC hotspot stimulation.** Table shows counts of neurons expressing Fos+ protein in various mesocorticolimbic structures and subregions after final exposure to rostromedial OFC hotspot laser stimulation in ChR2 rats (N = 7), eYFP controls (N = 5), and naïve rats (N = 6). Fos+ counts reflect mean of each group at each site ± standard error (SEM). One-way ANOVA's or Kruskal-Wallis was performed followed by corrected, two sided-post hoc tests between ChR2 and eYFP or ChR2 and naïve rats. *$p < 0.05$, **$p < 0.01$, ***$p < 0.001$, ****$p < 0.0001$.

| Insula Hotspot Fos + Neurons | Fos+ Counts (MEAN ±SEM) | | | Statistic | | Adjusted $p$ vs eYFP | 95% CI | Effect Size |
|---|---|---|---|---|---|---|---|---|
| | ChR2 | eYFP | Naïve | | | | | |
| Brain Region | n = 5 | n = 8 | n = 6 | *F (ANOVA) or H[#] (Kruskal- Wallis)* | *p* | *p* | | *d or $\eta^{2\#}$* |
| Rostromedial OFC | 273.6 ± 42.7 | 131.5 ± 20.7 | 101.0 ± 16.7 | 11.42[#] | 0.0006*** | 0.02* | | 0.59[#] |
| Caudal OFC | 103.6 ± 7.2 | 71.1 ± 14.3 | 42.7 ± 9.1 | 5.4 | 0.02* | 0.16 | | |
| Prelimbic Cortex | 125.4 ± 12.14 | 65.63 ± 11.25 | 59.0 ±8.9 | 9.43 | 0.002** | 0.003** | (20.6, 99.0) | 2.0 |
| Infralimbic Cortex | 81.00 ± 6.4 | 56.9 ± 6.1 | 53.3 ± 5.6 | 5.12 | 0.02* | 0.03* | (2.3, 46.0) | 1.5 |
| Rostral Insula | 36.2 ± 11.9 | 45.6 ± 6.5 | 46.5 ± 4.5 | 0.49 | 0.62 | | | |
| Rostral Anterior Cingulate | 68.2 ± 9.5 | 49.0 ± 8.4 | 31.8 ± 1.9 | 4.96 | 0.02* | 0.2 | | |
| Mid Anterior Cingulate | 71.8 ± 12.9 | 25.5 ± 6.8 | 16.8 ± 5.4 | 11.23 | 0.0009*** | 0.002** | (17.4, 75.3) | 2.1 |
| NAc Core | 53.6 ± 8.8 | 29.6 ± 2.8 | 30.5 ± 3.1 | 7.23 | 0.006** | 0.006** | (7.2, 40.8) | 1.6 |
| NAc Dorsal Medial Shell | 98.8 ± 5.9 | 50.1 ± 10.9 | 41.5 ± 4.8 | 8.39[#] | 0.009** | 0.03* | | 0.40[#] |
| NAc Ventral Medial Shell | 37.4 ± 8.0 | 36.0 ± 8.9 | 23.3 ± 3.6 | 0.95 | 0.41 | | | |
| NAc Caudal Dorsal Medial Shell | 60.8 ± 6.9 | 34.6 ± 5.5 | 36.7 ± 4.1 | 6.06 | 0.01* | 0.009** | (6.5, 45.9) | 1.7 |
| Dorsolateral Striatum | 12.8 ± 1.8 | 19.4 ± 3.8 | 12.7 ± 1.6 | 1.71 | 0.21 | | | |
| Olfactory Tubercle | 33.2 ± 3.5 | 19.5 ± 3.0 | 13.8 ± 0.9 | 10.94 | 0.001* | 0.007* | (3.8, 23.6) | 1.6 |
| Anterior VP Hotspot | 16.2 ± 3.1 | 22.9 ± 3.5 | 17.2 ± 3.4 | 1.2 | 0.34 | | | |
| Posterior VP Hotspot | 26.2 ± 9.2 | 17.8 ± 3.1 | 14.3 ± 1.5 | 0.75 | 0.49 | | | |
| Bed Nucleus Stria Terminalis | 26.6 ± 3.1 | 28.3 ± 4.0 | 26.5 ± 3.0 | 0.08 | 0.92 | | | |
| Lateral Hypothalamus | 58.6 ± 7.3 | 28.9 ± 4.3 | 20.8 ± 2.1 | 15.29 | 0.0002*** | 0.0009*** | (13.1, 46.4) | 2.1 |
| Perifornical Area | 25.2 ± 7.6 | 29.4 ± 4.4 | 16.0 ± 2.7 | 2.03 | 0.16 | | | |
| Central Amygdala | 24.2 ± 5.2 | 15.3 ± 3.3 | 15.2 ± 2.5 | 1.77 | 0.2 | | | |
| Basolateral Amygdala | 32.0 ± 8.6 | 18.4 ± 3.2 | 27.8 ± 4.5 | 1.92 | 0.19 | | | |
| Medial Amygdala | 58.8± 4.7 | 23.0 ± 3.6 | 23.0 ± 2.6 | 27.68 | <0.0001**** | <0.0001**** | (22.8, 48.8) | 3.5 |
| Paraventricular Thalamus | 34.4 ± 4.7 | 17.3 ± 4.2 | 14.5 ± 3.2 | 8.33[#] | 0.009** | 0.02* | | 0.40 |
| Arcuate Nucleus | 36.6 ± 8.3 | 16.3 ± 2.2 | 16.8 ± 2.0 | 6.77 | 0.007** | 0.007** | (5.6, 35.1) | 1.5 |
| Substantia Nigra | 21.6 ± 10.3 | 9.3 ± 2.0 | 7.3 ± 3.2 | 1.98 | 0.17 | | | |
| Ventral Tegmental Area | 62.6 ± 10.9 | 29.1 ± 3.0 | 16.5 ± 2.3 | 16.12 | 0.0005*** | 0.001** | (14.1, 52.9) | 1.78 |
| Periacqueductal Gray | 20.2 ± 4.3 | 15.3 ± 2.7 | 11.8 ± 0.8 | 1.93 | 0.18 | | | |

### Table S2.

**Raw Fos counts after caudal insula hotspot stimulation.** Table shows counts of neurons expressing Fos+ protein in various mesocorticolimbic structures and subregions final exposure to caudal insula hotspot laser stimulation in ChR2 rats (N = 5), eYFP controls (N = 8), and naïve rats (N = 6). Fos+ counts reflect mean of each group at each site ± standard error (SEM). One-way ANOVA's or Kruskal-Wallis was performed followed by corrected, two sided-post hoc tests between ChR2 and eYFP or ChR2 and naïve rats. *$p < 0.05$, **$p < 0.01$, ***$p < 0.001$, ****$p < 0.0001$.

| OFC/Insula Coldstrip Fos+ Neurons | Fos+ Counts (MEAN ±SEM) | | | Statistic | | Adjusted *p* vs eYFP | 95% CI | Effect Size |
|---|---|---|---|---|---|---|---|---|
| | ChR2 | eYFP | Naïve | | | | | |
| Brain Region | n = 5 Insula, n = 7 OFC | n = 6 | n = 6 | *F (ANOVA) or H# (Kruskal- Wallis)* | *p* | *p* | | *d or* $\eta^{2\#}$ |
| Rostromedial OFC | 93.9 ± 14.8 | 110.8 ± 30.3 | 101.0 ± 16.7 | 0.19 | 0.83 | | | |
| Caudal OFC *Counts from Insula ChR2 rats only | 136.6 ± 18.4 | 53.5 ± 8.7 | 42.7 ± 9.1 | 16.79 | 0.0002*** | 0.0006*** | (38.9, 126.1) | 2.5 |
| Prelimbic Cortex | 108.5 ± 11.3 | 50.0 ± 3.4 | 59.0 ±8.9 | 15.25# | 0.0005*** | 0.0011 | | 0.63# |
| Infralimbic Cortex | 99.4 ± 10.1 | 42.5 ± 5.8 | 53.3 ± 5.6 | 10.47 | 0.0006*** | 0.001** | (23.2, 90.6) | 2.1 |
| Rostral Insula **Counts from OFC ChR2 rats only | 87.6 ± 8.9 | 50.0 ± 10.6 | 46.5 ± 4.5 | 7.56 | 0.005** | 0.01* | (8.1, 67.2) | 1.5 |
| Caudal Insula | 66.2 ± 14.3 | 52.5 ± 13.8 | 42.2 ± 7.2 | 2.50# | 0.3 | | | |
| Rostral Anterior Cingulate | 64.3 ± 8.5 | 37.2 ± 8.3 | 34.5 ± 5.2 | 4.14 | 0.04* | 0.07 | | |
| Mid Anterior Cingulate | 18.4 ± 4.1 | 18.8 ± 10.9 | 24.2 ± 6.7 | 0.21 | 0.81 | | | |
| NAc Core | 62.8 ± 7.3 | 40.3 ± 3.7 | 30.5 ± 3.1 | 6.54 | 0.006** | 0.06 | | |
| NAc Rostrodorsal Medial Shell | 30.8 ± 6.2 | 38.5 ± 11.8 | 41.5 ± 4.8 | 0.56 | 0.58 | | | |
| NAc Rostroventral Medial Shell | 19.0 ± 4.5 | 19.7 ± 7.4 | 23.3 ± 3.6 | 1.53# | 0.47 | | | |
| NAc Caudal Dorsal Medial Shell | 71.1 ± 6.1 | 36.0 ± 2.0 | 36.7 ± 4.1 | 15.45# | 0.0004*** | 0.002** | | 0.64# |
| Dorsolateral Striatum | 7.6 ± 1.5 | 15.33 ± 6.7 | 12.7 ± 1.6 | 3.51# | 0.17 | | | |
| Olfactory Tubercle | 15.5 ± 2.8 | 14.0 ± 1.9 | 13.8 ± 0.9 | 0.12# | 0.94 | | | |
| Anterior VP Hotspot | 51.8 ± 3.6 | 30.7 ± 1.8 | 17.2 ± 3.4 | 17.48 | <0.0001**** | 0.0008*** | (8.9, 33.4) | 2.32 |
| Posterior VP Hotspot | 8.6 ± 3.9 | 5.7 ± 1.2 | 14.3 ± 1.5 | 8.1# | 0.02* | >0.9999 | | |
| Bed Nucleus Stria Terminalis | 18.4 ± 4.1 | 18.8 ± 10.9 | 26.5 ± 3.0 | 3.98# | 0.14 | | | |
| Lateral Hypothalamus | 38.9 ± 9.0 | 15.5 ± 4.7 | 20.8 ± 2.1 | 3.24# | 0.20 | | | |
| Perifornical Area | 23.0 ± 4.9 | 28.3 ± 8.42 | 16.0 ± 2.7 | 0.87 | 0.43 | | | |
| Central Amygdala | 40.7 ± 7.8 | 13.7 ± 2.8 | 15.2 ± 2.5 | 10.52# | 0.005** | 0.008** | | 0.41# |
| Basolateral Amygdala | 53.8 ± 12.8 | 17.3 ± 3.4 | 27.8 ± 4.5 | 8.91# | 0.012* | 0.006** | | 0.33# |
| Medial Amygdala | 113.3 ± 16.8 | 52.0 ± 11.0 | 30.7 ± 3.7 | 14.52# | 0.0007*** | 0.06 | | |
| Paraventricular Thalamus | 61.6 ± 7.1 | 21.3 ± 7.5 | 16.2 ± 3.0 | 13.46 | 0.0002*** | 0.001** | (15.8, 64.7) | 1.90 |
| Arcuate Nucleus | 53.9 ± 7.8 | 17.7 ± 5.4 | 20.2 ± 4.1 | 13.75# | 0.001** | 0.003** | | 0.56# |
| Substantia Nigra | 7.2 ± 1.4 | 8.0 ± 2.1 | 7.3 ± 3.2 | 0.03 | 0.97 | | | |
| Ventral Tegmental Area | 36.2 ± 6.6 | 13.5 ± 3.3 | 16.5 ± 2.3 | 5.85 | 0.01* | 0.01** | (4.5, 41.0) | 1.51 |
| Periacqueductal Gray | 17.3 ± 2.8 | 28.7 ± 3.8 | 11.8 ± 0.8 | 7.9 | 0.003** | 0.02* | (-21.0, -1.6) | -1.28 |

### Table S3.

**Raw Fos counts after caudal OFC/ rostral insula coldstrip stimulation.** Table shows counts of neurons expressing Fos+ protein in various mesocorticolimbic structures and subregions after final exposure to caudal insula hotspot laser stimulation in ChR2 rats (N = 7 caudal OFC; N = 5 rostral insula), eYFP controls (N = 6), and naïve rats (N = 6). Fos+ counts reflect mean of each group at each site ± standard error (SEM). One-way ANOVA's or Kruskal-Wallis was performed followed by corrected, two sided-post hoc tests between ChR2 and eYFP or ChR2 and naïve rats. *$p < 0.05$, **$p < 0.01$, ***$p < 0.001$, ****$p < 0.0001$.
